# Network Statistics of the Whole-Brain Connectome of *Drosophila*

**DOI:** 10.1101/2023.07.29.551086

**Authors:** Albert Lin, Runzhe Yang, Sven Dorkenwald, Arie Matsliah, Amy R. Sterling, Philipp Schlegel, Szi-chieh Yu, Claire E. McKellar, Marta Costa, Katharina Eichler, Alexander Shakeel Bates, Nils Eckstein, Jan Funke, Gregory S.X.E. Jefferis, Mala Murthy

## Abstract

Brains comprise complex networks of neurons and connections. Network analysis applied to the wiring diagrams of brains can offer insights into how brains support computations and regulate information flow. The completion of the first whole-brain connectome of an adult *Drosophila*, the largest connectome to date, containing 130,000 neurons and millions of connections, offers an unprecedented opportunity to analyze its network properties and topological features. To gain insights into local connectivity, we computed the prevalence of two- and three-node network motifs, examined their strengths and neurotransmitter compositions, and compared these topological metrics with wiring diagrams of other animals. We discovered that the network of the fly brain displays rich club organization, with a large population (30% percent of the connectome) of highly connected neurons. We identified subsets of rich club neurons that may serve as integrators or broadcasters of signals. Finally, we examined subnetworks based on 78 anatomically defined brain regions or neuropils. These data products are shared within the FlyWire Codex and will serve as a foundation for models and experiments exploring the relationship between neural activity and anatomical structure.

## Introduction

Mathematical network theory has been applied to connectomes at multiple scales (from detailed synaptic-resolution wiring diagrams to putative connectivity between brain regions) to understand brainwide organization (1–7). Network analyses quantify the interconnectivity and robustness of a network(8–10), and can identify highly connected nodes in the brain that may act as hubs (11). Such analyses can also serve as a basis for comparison across brain regions, individuals, developmental stages, or species, enabling researchers to uncover commonalities and differences in brain organization.

Mesoscale connectomes have been constructed for the brains of humans and other mammals from, for example, MRI and MEG data, which assess connectivity at millimeter scale (1, 12–15), relying on functional correlations in activity to infer mesoscale connectivity. Rich club organization has been observed in several mesoscale connectomes, including *Drosophila* (16, 17), humans, and other mammals (3, 4, 14). It has been suggested that such a network architecture contributes to the ability of brains to efficiently integrate and disseminate information.

Advancements in electron microscopy and dense volumetric reconstruction have enabled researchers to examine increasingly larger brain networks at the microscale. These methods do not make assumptions about the relationship between neuron connectivity and functional correlations. In network analyses performed at the microscale, nodes and edges can be directly related to neurons and synaptic connections. For instance, in the rich club regime observed in the *C. elegans* connectome, many rich club neurons are known to be important in motor control (2, 18, 19). Recurring patterns of connectivity between neurons, known as network motifs, have been proposed as “building blocks” of networks (20, 21), and their prevalence in neuronal networks has been studied to uncover organizational principles of neural networks (2, 5–7, 22–24). Specific motifs such as reciprocal connections (2, 6, 7, 19, 25), feedforward loops (2, 22, 23), and 3-unicycles (7, 26) have received significant attention in neuroscience because of their implications for local computation and information flow.

In this study, we characterize the network properties of the FlyWire synapse-resolution connectome, the first complete wiring diagram of an adult fly brain (27–30). We explore the interconnectivity of the brain, including path lengths between neurons, frequently traversed neural sub-populations, motif frequencies, and more. We draw statistical comparisons between the network of the fly brain and other biological wiring diagrams. We find that the fly brain has rich club organization and examine several sub-populations of these well-connected neurons, including those which may act as integrators or broadcasters of signals. Finally, we uncover differences in connectivity between 78 anatomically defined brain regions. The data derived in this work offer a quantitative summary of the network of the adult fly brain, and lay the groundwork for future studies exploring connectivity in the fly. They also serve as a valuable foundation for future experimental and theoretical work. A summary of computed statistics and neuron populations can be found in **Table 1**.

**Table 1.**
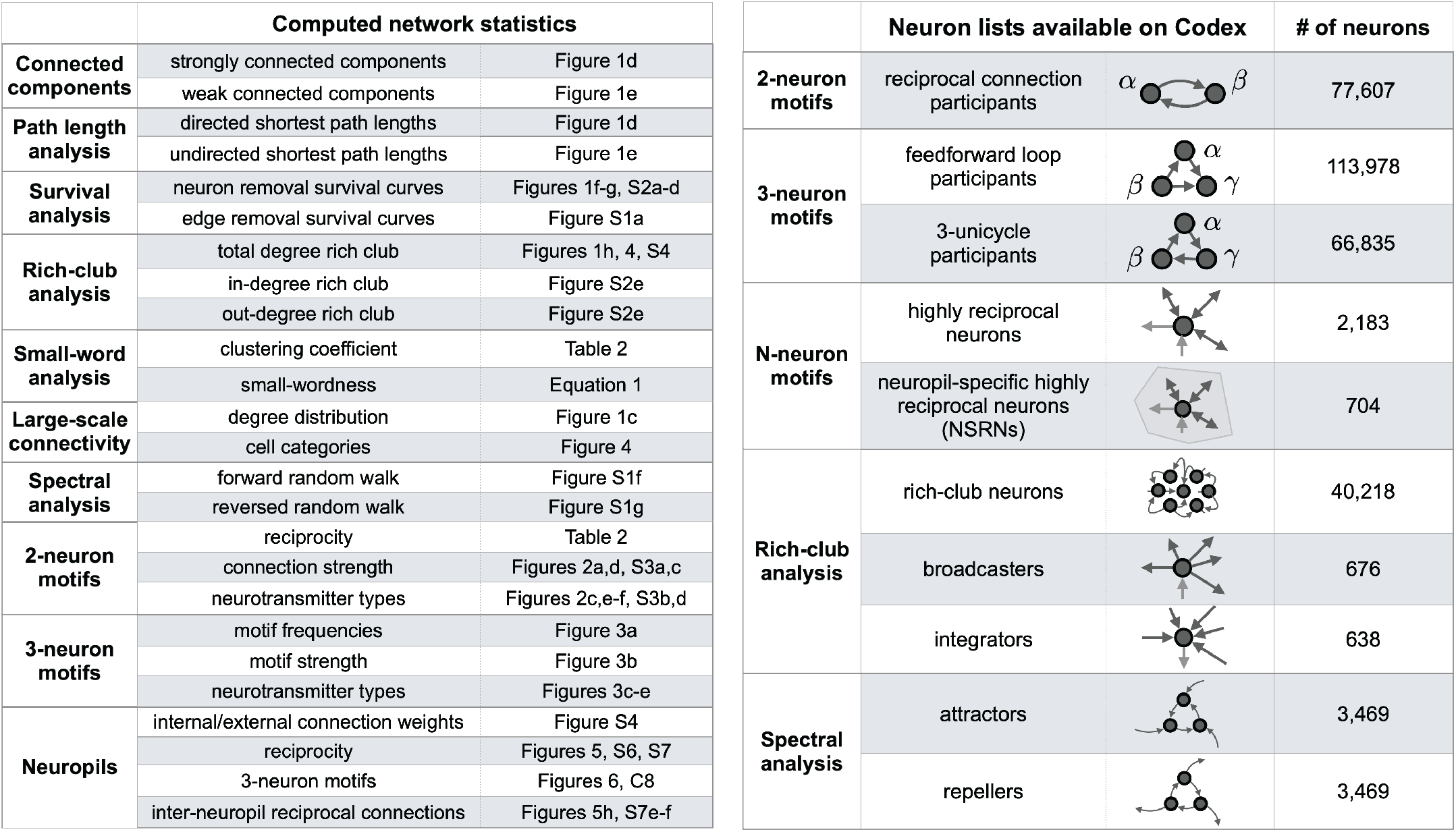
Data availability. List of data products in this work, including statistics computed in this paper (left) and neuron populations (right). Complete, interactive neuron lists are available online as “Connectivity Tags” on Codex (`codex.flywire.ai`). Definitions for each of these neuron populations can be found in the text, and in **Table S1**.

## Results

### Summary of the dataset and definitions

To perform large-scale network analyses, we summarized the synaptic connections between neurons into the following data structure. For each pair of neurons, we sum the total number of synaptic connections to return the weight of their connection. Repeating for all neuron pairs gives us a weighted graph describing the connectome, with 127,978 neurons and 2,613,129 total thresholded connections, representing the complete *Drosophila* brain (28) (**Figure 1a, Methods**). In this paper, we will be using the term “connection” to denote an edge that exists in the network between two neurons, consisting of one or more synapses. The synapses in this dataset were detected automatically (31, 32). To minimize the impact of spurious synapses, we applied a threshold of 5 synapses per connection for all of the analyses conducted in this study, unless otherwise noted (**Methods**). The exceptions are the distribution of synapses per connection, which is presented without threshold (**Figure 1b**), and controls to confirm that our qualitative observations are robust to threshold choice (**Figure S1b-c, Table S2**). We will be using synapse count as a proxy for edge strength in this paper: “stronger” and “weaker” will refer to higher or lower synapse counts, respectively.

**Figure 1.**
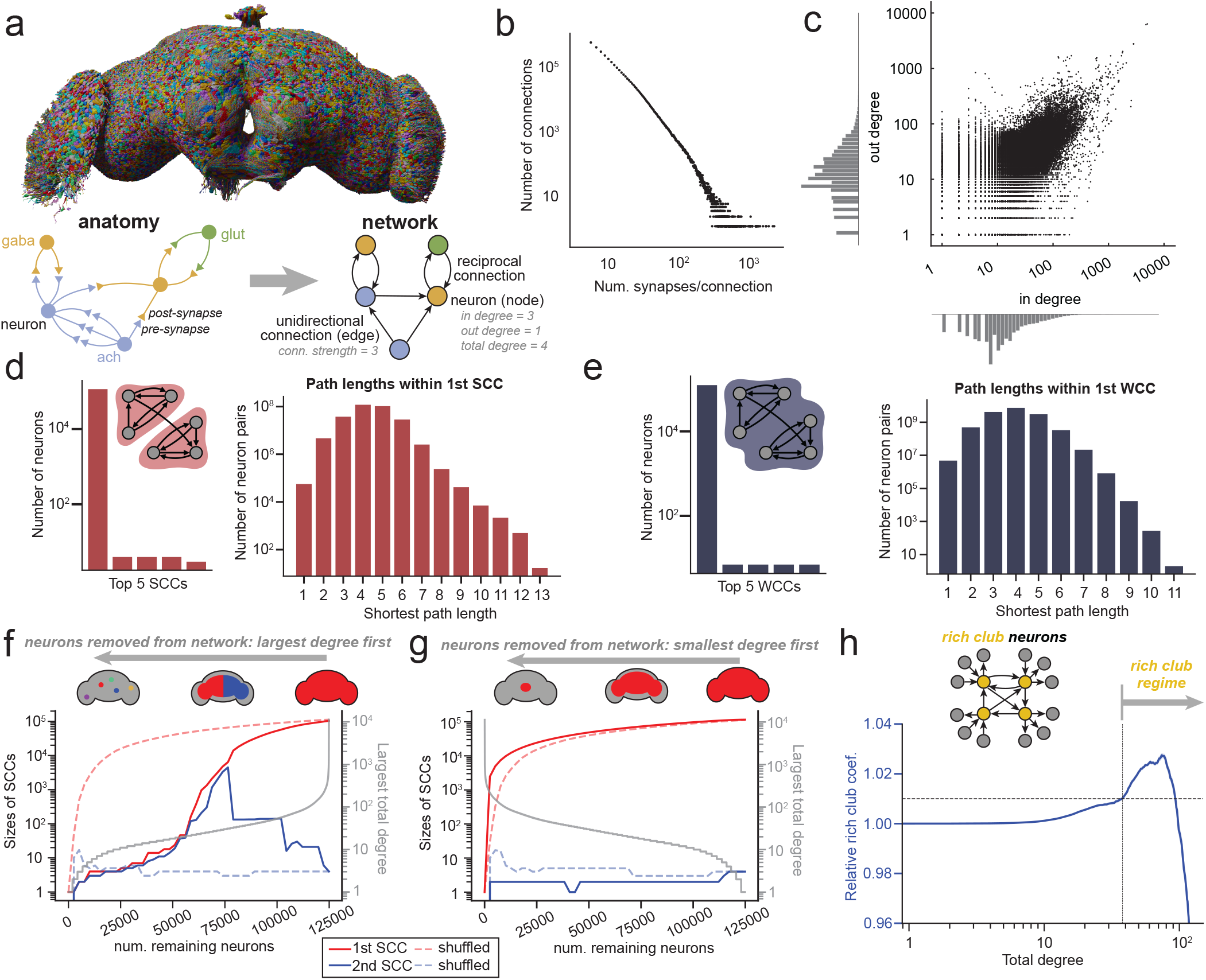
Whole-brain network properties. **(a)** The FlyWire dataset (27, 28, 30) is an EM reconstruction of the complete brain of an adult female *Drosophila melanogaster*, with both hemispheres of the brain and both optic lobes. The volume contains 127,978 neurons and 32 million synapses with a threshold of 5 synapses/connection applied (subsample of synapse locations shown in the inset). **(b)** The distribution of the number of synapses per connected neuron pair. **(c)** The in-degree (number of presynaptic partners) plotted against the out-degree (number of post-synaptic partners), with log-scale x and y-axes. **(d)** Strongly connected components (SCCs) consist of a subset of nodes in a network which are mutually reachable via directed edges. In the fly brain there exists one giant SCC containing 93.3% of all neurons after thresholding at 5 synapses per connection. The distribution of shortest path lengths between neuron pairs within this SCC is plotted. **(e)** Weakly connected components (WCCs) consist of a subset of nodes in a network which are mutually reachable, regardless of edge direction. In the fly brain there exists one giant WCC containing 98.8% of all neurons. The distribution of path lengths between neuron pairs within this WCC is plotted. **(f)** We examine the role high-degree neurons play in connecting the brain by plotting the sizes of the first two strongly connected components (SCCs) as nodes are removed by total degree (2500 neurons per step). Removal of neurons starting with those with largest degree results in the brain splitting into two SCCs when neurons of approximately degree 50 start to be removed, a deviation from when neurons are removed in a random order (dotted lines). The largest surviving total degree as a function of the number of remaining nodes is plotted in gray. **(g)** Removal of neurons starting with those with smallest degree results in a single giant SCC until all neurons are removed (2500 neurons per step). The smallest surviving total degree as a function of the number of remaining nodes is plotted in gray. **(h)** The relative rich club coefficient as a function of total degree, computed relative to CFG null models. The range over which the relative rich club coefficient is greater than 1.01 is 37 to 93. We take all neurons with total degree > 37 to be within the rich club regime.

The FlyWire connectome also contains synapse-level neurotransmitter predictions (33). The classifier applied to the dataset discriminates between six neurotransmitters: the fast-acting classical neurotransmitters acetylcholine (ach), GABA (gaba), and glutamate (glut) and the monoamines dopamine (da), octopamine (oct), and seratonin (ser). In the *Drosophila* nervous system, acetylcholine is excitatory and GABA is inhibitory. Glutamate can be either excitatory or inhibitory, but within the brain of the fly it has largely been observed to be inhibitory (34–36).

A key characteristic of the network is the distribution of degrees, which reflects the amount of connectivity between neurons. For any given neuron, the *in-degree* is defined as the number of presynaptic neurons (neurons it receives inputs from), and the *out-degree* is defined as the number of postsynaptic neurons (neurons it sends outputs to). With a threshold of 5 synapses per connection, the average in/out-degree of an intrinsic neuron in the brain is 20.5 (28), but the distributions of in-degree and outdegree are not highly correlated (Pearson *R* = 0.76, *p <* 0.001) (**Figure 1c**). On average, each connection in the brain consists of approximately 12.6 synapses after the threshold is applied (28). Across the connectome, the probability that any two neurons is connected is 0.000161. This makes the wiring diagram of the fly brain a very sparse matrix when compared to, for example, the *C. elegans* nervous system or the partial wiring diagrams of brain regions of larval zebrafish and mouse (**Table 2**). This sparsity is due in part the size of the fly brain. The connection probability is highest among neurons whose arbors are close to each other. Over 71% of connections occur between neuron pairs located within 50 microns of each other, despite these pairs constituting less than 3% of the total number of pairs. (**Figure S1d**). We note, however, that even in the close regime the connection probability in the fly remains lower than what has been observed in other wiring diagrams. The longrange sparsity is partially a consequence of the segregation of the neurons of the *Drosophila* brain into a large number (78) of brain regions (neuropils), and we further investigate connectivity within neuropils below (**Figures 5-6, S5-S8**).

**Table 2.**
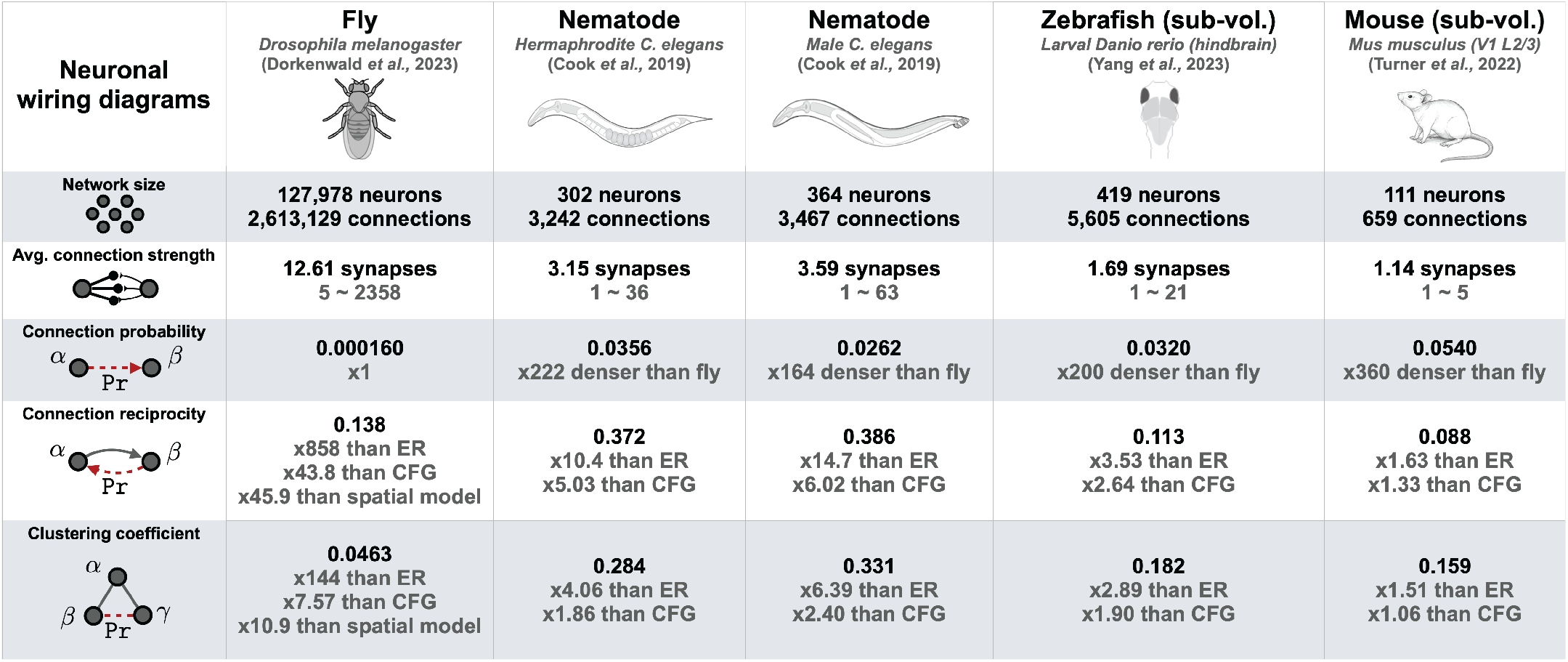
Connection probabilities, reciprocity, and clustering coefficient in the fly brain. The probability that any two neurons in the fly brain are connected is 0.000160. Connection reciprocity (the probability that two connected neurons are reciprocally connected) in the fly is 0.138, larger than in either an ER, CFG, or spatial null model **(Methods)** with the same sparsity. The clustering coefficient (the probability that if neuron *α* and neuron *β* are connected and neuron *α* and neuron *γ* are connected, then neuron *β* and neuron *γ* are also connected, irrespective of directionality) in the fly is 0.0463. Both reciprocity and clustering coefficient are higher than expected with ER, CFG, and spatial null models. Values for thresholds from 0 to 50 are plotted in **Figure S1c**. Statistics for *C. elegans* were computed for the chemical networks of neurons in hermaphrodite and male worms (19). Statistics for larval zebrafish hindbrain (7) and mouse visual cortex (6) were computed excluding any truncated neurons

| Neuronal wiring diagrams | Fly<br>*Drosophila melanogaster*<br>(Dorkenwald et al., 2023) | Nematode<br>*Hermaphrodite C. elegans*<br>(Cook et al., 2019) | Nematode<br>*Male C. elegans*<br>(Cook et al., 2019) | Zebrafish (sub-vol.)<br>*Larval Danio rerio (hindbrain)*<br>(Yang et al., 2023) | Mouse (sub-vol.)<br>*Mus musculus (V1 L2/3)*<br>(Turner et al., 2022) |
| --- | --- | --- | --- | --- | --- |
| Network size | 127,978 neurons<br>2,613,129 connections | 302 neurons<br>3,242 connections | 364 neurons<br>3,467 connections | 419 neurons<br>5,605 connections | 111 neurons<br>659 connections |
| Avg. connection strength | 12.61 synapses<br>5 ~ 2358 | 3.15 synapses<br>1 ~ 36 | 3.59 synapses<br>1 ~ 63 | 1.69 synapses<br>1 ~ 21 | 1.14 synapses<br>1 ~ 5 |
| Connection probability | 0.000160<br>x1 | 0.0356<br>x222 denser than fly | 0.0262<br>x164 denser than fly | 0.0320<br>x200 denser than fly | 0.0540<br>x360 denser than fly |
| Connection reciprocity | 0.138<br>x858 than ER<br>x43.8 than CFG<br>x45.9 than spatial model | 0.372<br>x10.4 than ER<br>x5.03 than CFG | 0.386<br>x14.7 than ER<br>x6.02 than CFG | 0.113<br>x3.53 than ER<br>x2.64 than CFG | 0.088<br>x1.63 than ER<br>x1.33 than CFG |
| Clustering coefficient | 0.0463<br>x144 than ER<br>x7.57 than CFG<br>x10.9 than spatial model | 0.284<br>x4.06 than ER<br>x1.86 than CFG | 0.331<br>x6.39 than ER<br>x2.40 than CFG | 0.182<br>x2.89 than ER<br>x1.90 than CFG | 0.159<br>x1.51 than ER<br>x1.06 than CFG |

### Neurons in the brain form a single connected component

To assess the interconnectivity of the neurons in the brain, we searched the connectome for connected components using two sets of criteria. First, we looked for *strongly connected components* (SCCs). All neurons within an SCC are mutually reachable via directed pathways (37). Second, we looked for *weakly connected components* (WCCs), a relaxed criterion in which all neurons within a WCC are mutually reachable, ignoring the directionality of connections.

Despite its sparsity, the brain is highly connected under either criteria – 93.3% of neurons are contained in a single SCC, while 98.8% of neurons are contained in a single WCC (**Figure 1d-e**). These giant connected components, which contain the overwhelming majority of neurons in the brain, persist when either the strongest connections or the weakest connections are pruned (**Figure S1a-b**), indicating that connectivity in the brain is robust: many paths connect neuron pairs. We will refer to these extremely large connected components as the giant SCC and giant WCC, respectively. Within the giant SCC, the average shortest directed path length between neuron pairs is 4.42 hops, with every neuron reachable within 13 hops (**Figure 1d**). In the giant WCC, the average shortest undirected path length between neuron pairs is 3.91 hops, with every neuron reachable within 11 hops (**Figure 1e**). These numbers are comparable to those found in a similar analysis of the hemibrain dataset (38). The short path lengths within both connected components show that despite its size, the fly brain is still relatively shallow when compared to artificial networks (39).

Is the high interconnectivity observed in the fly brain a consequence of a relatively large number of interconnected neurons, or is it dependent on a small number of very highly connected “hub” neurons? To assess this, we constructed survival curves, observing for how long the connected components of the network persist when neurons are removed from the network. Here, we plot the sizes of the two largest SCCs as we remove neurons from the directed network, starting with those of largest total degree (**Figure 1f**). We find that the first giant SCC persists until a total degree of 50, at which point the network splits into two SCCs of roughly equal size. These two SCCs correspond to a split between the left and right hemispheres, and demonstrate that despite the hemispheric anatomy of the brain, the two hemispheres are highly interconnected: they do not split into separate networks until about 60% of all neurons are removed. Removing neurons from the network by smallest total degree does not result in division of the first giant connected component (**Figure 1g**). This indicates that the interconnectivity of the brain is robust, and not dependent on a small number of highly connected neurons. We observe similar behavior in the WCCs when removing neurons from the undirected network (**Figure S2a-b**). These results also remain qualitatively consistent when neurons are pruned in either the directed or undirected network by either in-degree or out-degree alone (**Figure S2c-d**).

The SCC criteria is more biologically realistic, since connections between real neurons are directed. Note, however, that the similarities in size and path length distribution between the first SCC and first WCC indicate the prevalence of recurrent connections in the brain. In a mostly feedforward network, one would expect a smaller SCC with longer path lengths. This is not what we observe in the fly brain—instead, across the population of all neuron pairs, the distribution of shortest directed path lengths is comparable to the distribution of shortest undirected path lengths.

### Spectral analysis of the whole-brain network

To better understand the network topology of the brain, we performed a spectral analysis of a random walk in the giant SCC. In this random walk, the transition probability from neuron *α* to neuron *β* is 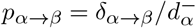, where 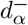 is the out-degree of neuron *α*, and *δ*_*α→β*_ *∈ {*0, 1*}* indicates the existence of a connection. Such a random walk converges to a stationary distribution over all neurons in the giant SCC (**Figure S1f**). We found that in this random walk, 3% of neurons were visited 61.2% of the time—the remaining 97% of neurons were visited only 38.8% of the time. These top visited neurons can therefore be classified as *attractor nodes* (40) in the network. These attractor nodes typically make connections in the gnathal ganglia (GNG), a large midline neuropil which both sends and receives information from the periphery and contains a large number of neurons that connect to the ventral nerve cord (VNC).

We also performed a “reverse” walk within the giant SCC, reversing edge directionality so that the transition probability from neuron *α* to neuron *β* is 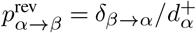, where 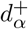 is the in-degree of neuron *α*. The reversed walk also converges to a stationary distribution in which 3% of neurons were visited 42.4% of the time (**Figure S1g**). These highly visited neurons in a reverse random walk are *repeller nodes* in the network. Many of these neurons make synapses in the antennal lobes (AL) and medullae (ME), brain regions close to the olfactory and visual periphery, respectively. This suggests that these neuropils engage in local (rather than integrative) computations.

### The fly brain has a large rich club

Many networks exhibit the “rich club” property (3, 11, 18), in which well connected nodes are preferentially connected to other well connected nodes (see **Methods**). We find that there exists a rich club regime in the FlyWire connectome, in which neurons are more highly interconnected than one would expect from a randomly connected network (**Figure 1h**). We will take this cutoff to be a total degree of 37, though we note that the exact choice of rich club cutoff is arbitrary (**Methods**). This large rich club regime contains 40,218 neurons, approximately 30% of all neurons in the brain. The connection probability within this rich club is 0.000870, 5.4 times higher than the overall connection probability in the brain. Such a large rich club suggests that the topology of the fly wiring diagram is fairly distributed. This is consistent with the connected component observations, which also suggest a degree of robustness. A rich club analysis considering in-degree alone returns an in-degree threshold of 10, while no rich club is observed when considering out-degree alone (**Figure S2e**).

The fraction of neurons in the rich club regime in the fly is substantially larger in the fly than in *C. elegans*, which has a rich club of 11 neurons (4% of the neurons in the worm) (18). We caution that this difference in rich club size is sensitive to the criteria used to determine the rich club cutoff, and may also be a consequence of the different scales of these two networks. Nonetheless, it is interesting to note that while the worm rich club contains known hub neurons, such as the command neurons AVA and AVB, such highly connected hub neurons do not seem to be present in the fly brain—while there are neurons with very high degrees, there also exist alternate paths between most neuron pairs. We further examine the properties of this large rich club population in the section: **Large-scale connectivity in the brain**.

### Reciprocal and recurrent motifs are over-represented in the brain

Connection reciprocity is a measure of the amount of direct feedback in the brain: given that neuron *α* is connected to neuron *β*, what is the probability that neuron *β* is connected back to neuron *α*? Across the whole brain, this connection reciprocity probability is 0.138 (**Table 2**). The connection reciprocity in the brain is significantly higher than in both the Erdős-Rényi (ER) and configuration (CFG) random null models (**Methods**). The over-representation of reciprocal connections in brains relative to null models is well established, and our results are consistent with previous observations both in *Drosophila* (38, 41, 42) and in other species (2, 6, 19, 22, 23, 43).

We also computed the clustering coefficient, a higher-order connectivity metric which assesses the prevalence of triplet structures in the network irrespective of edge direction: if neuron *α* and neuron *β* are connected and neuron *α* and neuron *γ* are connected, what is the probability that neuron *β* and neuron *γ* are also connected? The clustering coefficient in the brain is 0.0477 (**Table 2**). As was the case with reciprocity, this value of clustering coefficient is higher than in both ER and CFG null models. The high clustering coefficient demonstrates that the network of the fly brain is highly connected and is nonrandom in its structure.

We compared these metrics with two existing whole-animal connectomes, the hermaphrodite and male *C. elegans* (2, 19, 44), and with two sub-volume wiring diagrams, the hindbrain of a larval zebrafish (7) and a region of L2/3 mouse visual cortex (6) (**Table 2**). Despite differences in sparsity of the different brain networks, the values of reciprocity and clustering coefficient are comparable across all five datasets.

The fly brain is physically much larger than other previously studied biological networks, such as those in *C. elegans*, and it is divided into distinct brain regions. However, ER and CFG null models do not contain any spatial information, instead assuming that any neuron pairs may randomly connect. We therefore constructed a spatial null model to account for some of the physical constraints. Informed by the distribution of connections as a function of distance, we built a two-zone spatial null model, where the probability of randomly forming a connection between neurons is dependent on the distance between them (**Figure S1e**) (**Methods**). We computed the reciprocity and clustering coefficient for the spatial null model and found that reciprocity and clustering coefficient in the real network were also higher than this null model, suggesting that the non-random nature of connectivity in the fly is not solely a consequence of spatial or morphological constraints.

We note that interpretations of these direct comparisons of metrics across different datasets should be made with caution. While the fly and worm datasets represent complete brains and nervous systems, respectively, the zebrafish and mouse datasets are derived from brain sub-volumes, with order 100s of neurons. Because many neurons in the fish and mouse sub-volumes are truncated, measures of reciprocity and clustering coefficient are incomplete. Additionally, differences in synapse detection and synapse thresholding will impact topological metrics such as connection probability and reciprocity. While connectomes in *C. elegans* have been proofread to the level of individual synapses (2, 25, 44), it is not feasible to manually proofread every synapse in larger connectomics datasets such as *Drosophila*. Varying the synapse threshold in the fly did not significantly alter reciprocity and clustering coefficient values (**Figure S1c, Table S2**).

### Small-worldness of the fly brain

A “small-world” network is one in which nodes are highly clustered and path lengths are short (10). High small-worldness coefficients are associated with efficient communication between nodes (45, 46). We quantified the small-worldness of the connectome by comparing it to an Erdő s-Rényi (ER) graph (47). The average undirected path length in the ER graph, denoted as *ℓ*_rand_, is estimated to be 3.57 hops, similar to the observed average path length in the fly brain’s WCC (*ℓ*_obs_ = 3.91). The clus-tering coefficient (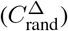) of the ER graph is only 0.0003, much smaller than the observed clustering coefficient (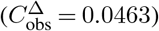) (**Table 2, Methods**). The small-worldness coefficient of the fly connectome is:

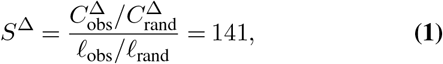

significantly higher than that of the *C. elegans* connectome (*S*^Δ^ = 3.21) and close to that of the internet (*S*^Δ^ = 98.1) (10), implying highly effective global communication among neurons in the brain.

### Strength and neurotransmitter composition of reciprocal connections

The average strength of edges participating in reciprocal connections is higher than the average strength of unidirectional connections (**Figure 2a**). The majority of unidirectional connections are cholinergic (excitatory), while edges participating in reciprocal connections contain fewer cholinergic neurons and more GABAergic neurons (**Figure 2b**). Inhibitory connections in the brain have more synapses on average than excitatory connections (28), which may partially explain the higher average strength of reciprocal connections. The most common reciprocal pairing is between a cholinergic neuron and a GABAergic neuron and the second most common pairing is acetylcholine-glutamate (**Figure 2c**). Both of these reciprocal motifs are excitatory-inhibitory (E-I), and both are over-represented when compared to the neurotransmitter frequencies observed for reciprocal connections (**Figure 2b**). Excitatory-excitatory (E-E) acetylcholine-acetylcholine pairs are in contrast under-represented, as are inhibitory-inhibitory (I-I) GABA-GABA pairs. We observed reciprocal E-I (acetylcholine-GABA and acetylcholine-glutamate) connection strengths to be only weakly correlated, while E-E (acetylcholine-acetylcholine) pairs were uncorrelated (**Figure 2d**). Examples of reciprocal neuron pairs are shown in **Figure 2g**.

**Figure 2.**
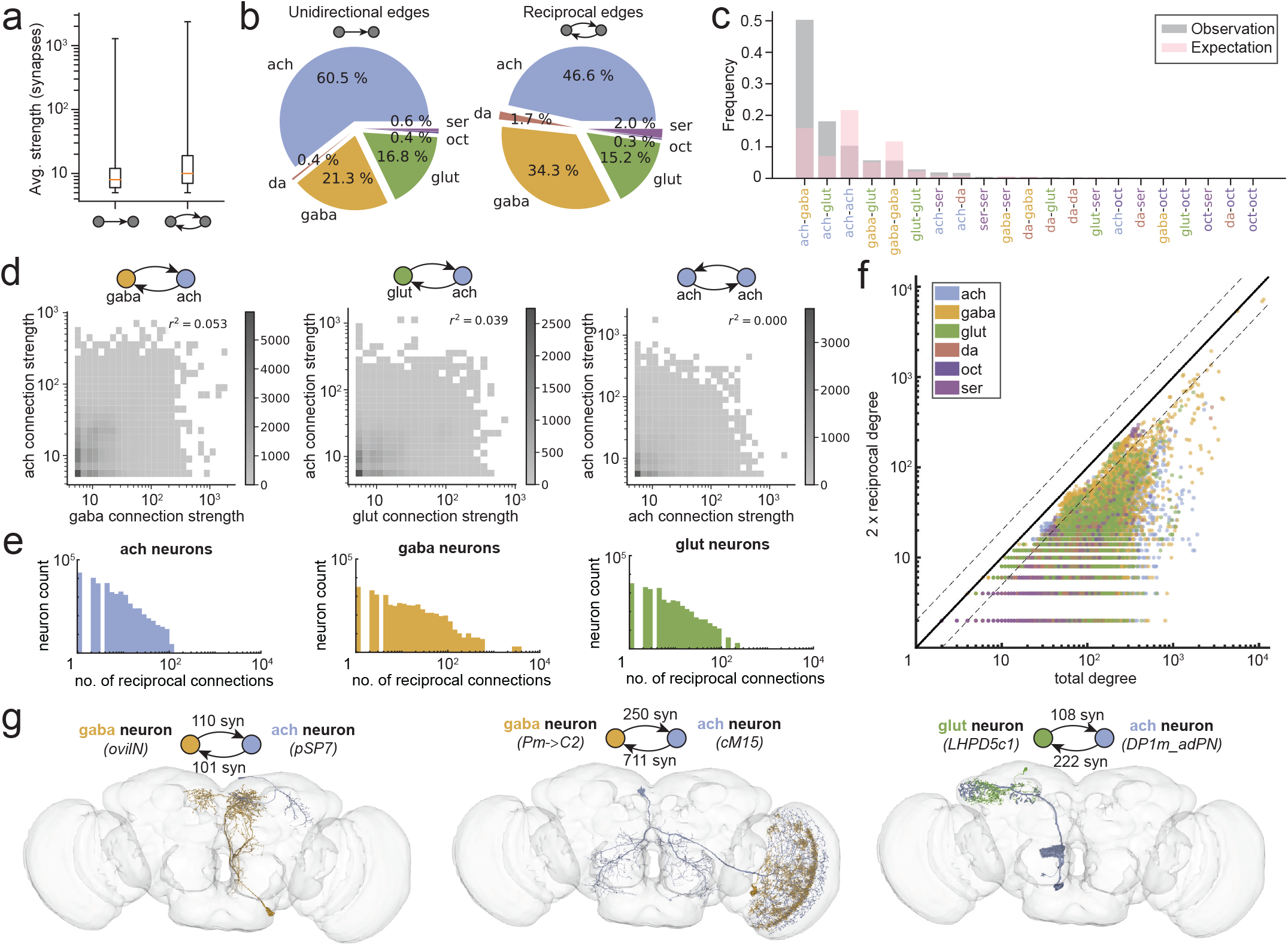
Characterizing reciprocal connections in the brain. **(a)** Edges that are part of reciprocal connections (reciprocal edges) are stronger on average than unidirectional connections. **(b)** Breakdown of unidirectional and reciprocal edges by neurotransmitter. Unidirectional connections are most likely to be cholinergic. Reciprocal connections are more likely than unidirectional connections to contain a GABAergic neuron. **(c)** The frequency of neurotransmitter pairs forming reciprocal connections, compared to the expected frequency of neurotransmitter pairs under the assumption of independent neurotransmitter choice (red). A majority of reciprocal connections are formed by acetylcholine-GABA pairs. The next most common reciprocal connection type is acetylcholine-glutamate, with acetylcholine-acetylcholine pairs under-represented. **(d)** Heatmaps of the relative strengths (synapse counts) of the two connections forming acetylcholine-GABA reciprocal pairs (left), acetylcholine-glutamate reciprocal pairs (center), and acetylcholine-acetylcholine reciprocal pairs (right). The strengths of the edges of reciprocal pairs are uncorrelated. Excitatory-inhibitory pairs (acetylcholine-GABA and acetylcholine-glutamate) have higher average strengths than excitatory-excitatory (acetylcholine-acetylcholine) pairs. **(e)** Distributions of reciprocal degree (the number of reciprocal connections a given neuron makes) for cholinergic neurons (left), GABAergic neurons (middle), and glutamatergic neurons (right). GABAergic neurons are more likely to make large numbers of reciprocal connections, while cholingeric neurons are more likely to have smaller numbers of reciprocal connections. **(f)** Scatterplot of 2 times the reciprocal degree of neurons versus their total degree (in-degree + out-degree). Dotted lines indicate a factor of 2 around the *x* = *y* line. Large neurons for which reciprocal connections form the majority of their total connections are most likely to be GABAergic. **(g)** Visualizations of exemplar reciprocal neuron pairs. Cell labels are listed where available.

### Reciprocal degree across the neuronal population

Of the 127,978 neurons in the whole brain, 77,607 participate in at least one reciprocal connection: approximately 2 in every 3 neurons, even with the synapse threshold we applied **(Methods)**. Many neurons participate in multiple reciprocal connections. To characterize these neurons, we define the *reciprocal degree* as the number of reciprocal connections made by a given neuron (**Figure S3a**). Plotting the distributions of reciprocal degree by neurotransmitter, we observe that the overwhelming majority of neurons with high reciprocal degree (*d*^rec^ *>* 100) are GABAergic (**Figures 2e, S3b**), while at lower reciprocal degrees (*d*^rec^ *<* 100), all three primary neurotransmitter types are well represented.

What fraction of a neuron’s connections are reciprocal? Note that here, we are not considering reciprocity between cell types, but rather between pairs of individual neurons. For most neurons these fractions are low—on average 23% of incoming and 18% of outgoing connections are reciprocal. Plotting the fraction of reciprocal incoming connections against the fraction of reciprocal outgoing connections, we observe only a weak correlation (**Figure S3c**), suggesting that a given neuron’s reciprocal degree is not strongly coupled to either its in-degree or its out-degree. Comparing the number of reciprocal connections neurons make to the total number of connections they make by plotting 2*×* the reciprocal degree against the total degree of neurons (in-degree + out-degree), we again see no relationship (**Figure 2f**). Dividing the neuron population by neurotransmitter, however, we find that neurons of high total degree are mostly GABAergic, and that for many of these neurons, more than half of their total connections are reciprocal (**Figure S3d**). Many of these highly reciprocal neurons provide feedback inhibition within specific neuropils (**Identifying neuropil-specific reciprocal neurons**). Examples of neurons which form reciprocal connections are shown in **Figure 2g**.

### Strength and neurotransmitter composition of three-node motifs

The high clustering coefficient of the brain implies an overrepresentation of triplet structures. We determined the frequency at which each of the 12 directed three-node motifs occur in the brain (**Figure 3a**). Feedforward motifs (motifs #1-3) are under-represented when compared to both ER and CFG null models, while all others, including the highly recurrent motifs (motifs #7-13), are over-represented. The strengths of edges participating in 3-node motifs are higher than the average edge strength (**Figure 3b**). Complex 3-node motifs which contain reciprocal connections tend to be stronger than feedforward motifs.

**Figure 3.**
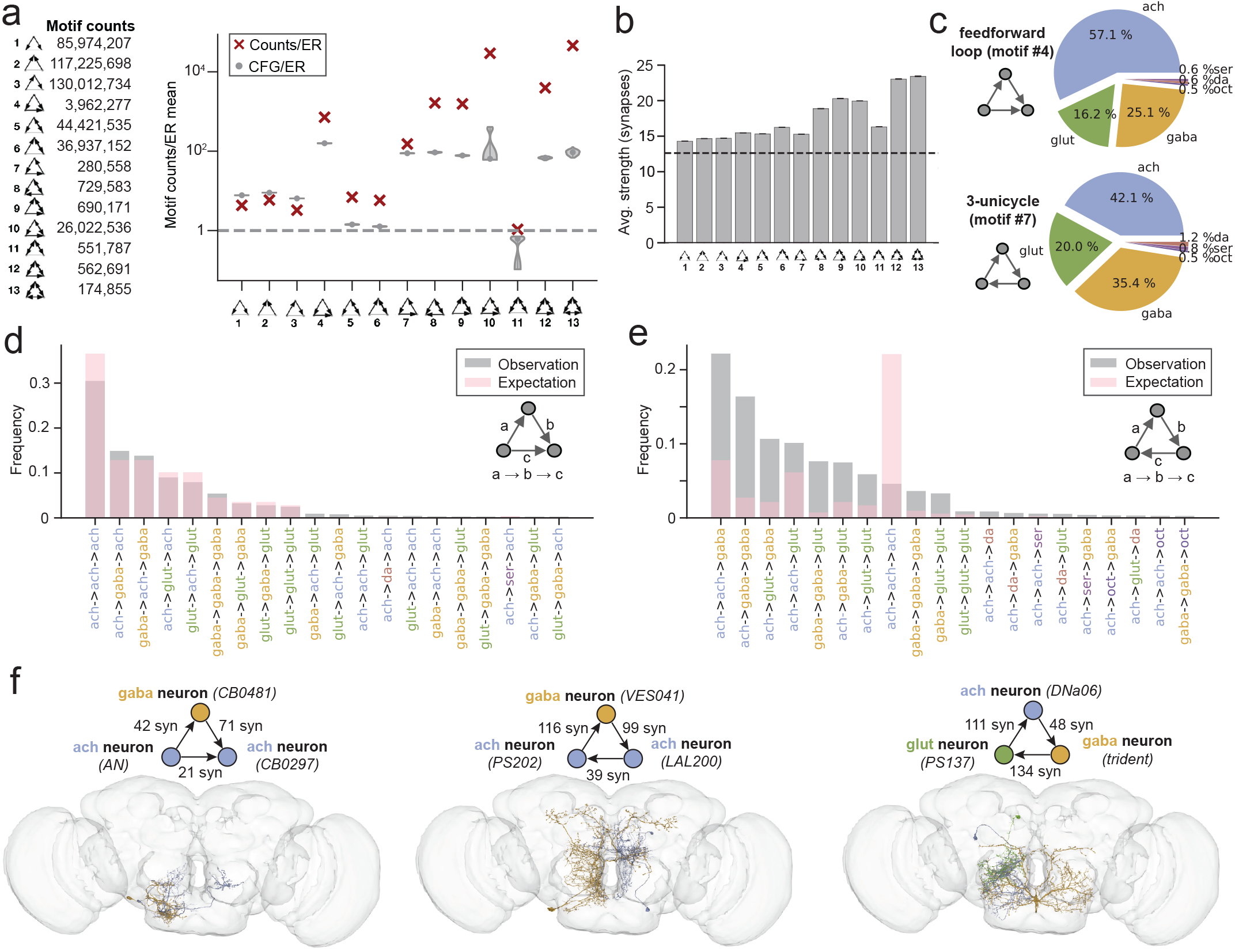
Examining 3-node motifs. **(a)** The distribution of three-node motifs across the whole brain. Absolute counts of each motif are on the left, and the frequency of each motif relative to that in an ER null model is plotted to the right, together with the average motif frequencies of 100 CFG models (gray violin plots). When we compare the whole-brain network to both ER and CFG null models, we observe an under-representation of simple motifs (#1-3) and an over-representation of other motifs, particularly highly recurrent motifs (#10, 12, 13). **(b)** The average strength of edges that are part of the 3-node motifs. The dotted line is the average connection strength in the brain. **(c)** Breakdown by neurotransmitter of edges participating in two motifs: feed-forward loops (motif #4) and 3-unicycles (motif #7). Edges in feed-forward loops are more likely to be cholinergic. **(d)** Further examining the neurotransmitter composition of these motifs, we find that feed-forward loops (motif #4) are most likely to be acetylcholine-acetylcholine-acetylcholine, **(e)** while 3-unicycles (motif #7) tend to contain at least one inhibitory edge (glutamate or GABA). **(f)** Visualizations of exemplar 3-node motifs. Cell labels are listed where available.

Examining the neurotransmitter composition of two of these three-node motifs, feedforward loops (motif #4) and 3-unicycles (motif #7) (**Figure 3c**), we found that edges which participate in feedforward loops were predominantly cholinergic, and that the most common neurotransmitter composition for a feedforward loop is three cholinergic neurons, a feed-forward excitatory configuration (**Figure 3d**). The next most common compositions contain either one or two inhibitory (GABAergic or glutamatergic) edges. Feedforward loops with one inhibitory edge are likely feedforward inhibition motifs, while loops with two inhibitory edges are likely disinhibition motifs. 3-unicycles in contrast contain a higher proportion of inhibitory GABAergic and glutamatergic neurons, and the three most common 3-unicycle compositions all contain at least one inhibitory neuron (**Figure 3e**). These cycles may act as indirect feedback inhibition circuits. It is interesting to note that the observed neurotransmitter composition frequencies are closer to what may be expected by chance for feedforward loops than they are for 3-unicycles. Examples of neurons which form 3-node motifs are shown in **Figure 3f**.

The fly brain exhibits a high clustering coefficient and an over-representation of highly connected 3-neuron motifs. These observations suggest that the local structure of the brain displays a high degree of non-randomness, in line with previous studies in *C. elegans* (2, 19) and in mouse cortex (6, 22, 23). The over-representation of feedforward loops (motif #4) has been widely observed in other biological networks, such as in rat cortex and *C. elegans* (2, 19, 22, 23). This over-representation is present in most neuropils in the brain. It is possible that these feed-forward loops, which are predominantly excitatory, may form large-scale feedforward structures which span brain regions. 3-unicycles (motif #7) may form recurrent local circuits capable of generating persistent oscillatory neural activity (7).

### Large-scale connectivity in the brain

Within the adult brain, the in-degree and out-degree of neurons are not tightly correlated. Neurons with few inputs and many outputs may serve as broadcasters of signals, while those with many inputs and few outputs may act as integrators. To examine these populations of neurons, we divided the intrinsic rich club neuron population into three categories based on their in-degree and out-degree (**Figure 4a**). We divided the rich club neurons by defining *broadcaster neurons* as those for which out-degree *≥* 5*×* in-degree, and *integrator neurons* as those for which in-degree *≥* 5*×* out-degree. The boundaries defining broadcaster and integrator neurons are arbitrary, and intended to aid in comparisons of neurons with unbalanced inputs and outputs. In the FlyWire connectome we find 676 broadcasters and 638 integrators. The remaining intrinsic rich club neurons (37,093) fall into the *balanced* category (Region 3), including most highly reciprocal neurons. Some examples of broadcasters, integrators, and balanced neurons are shown in **Figure 4d**.

**Figure 4.**
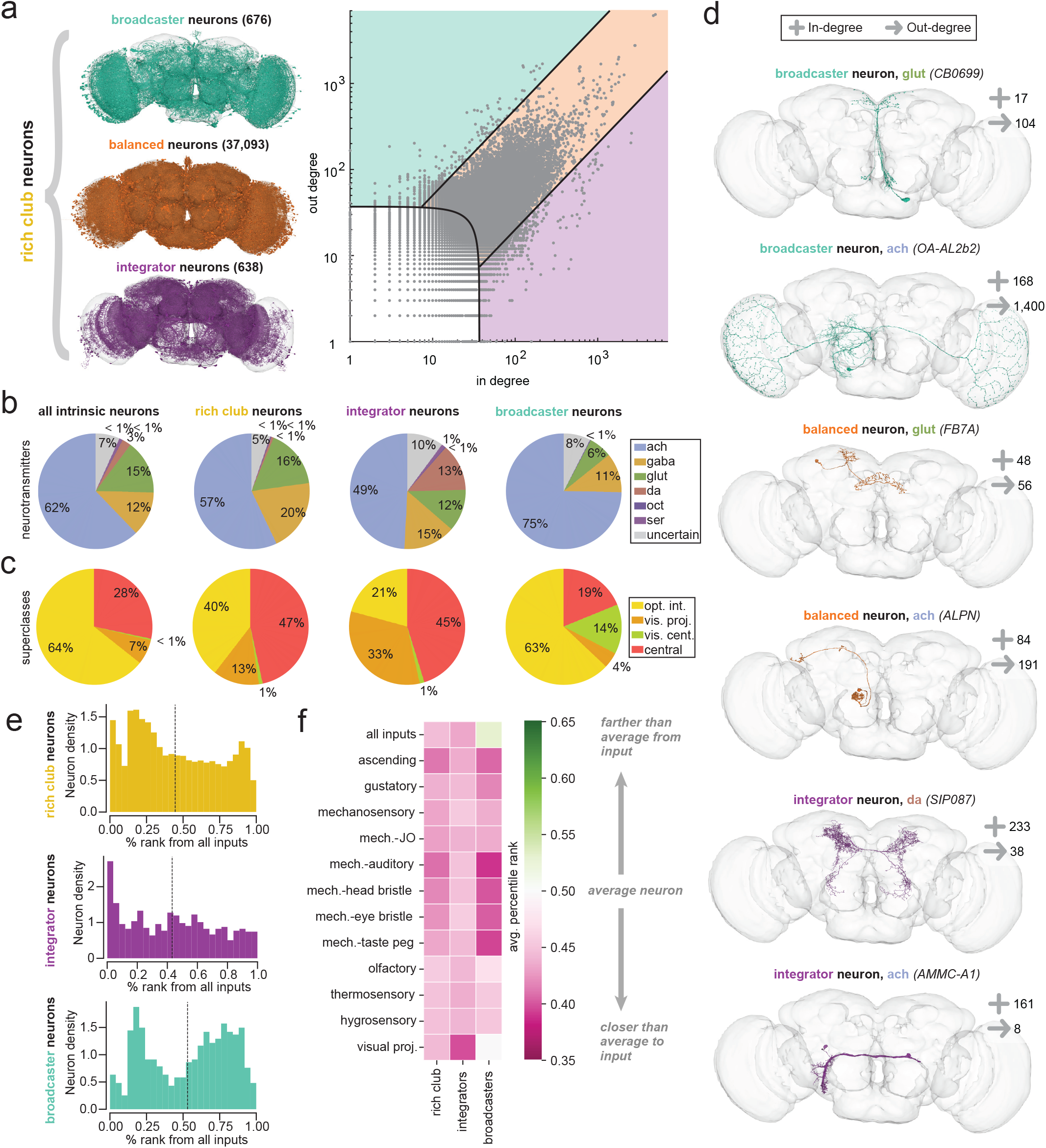
Large-scale neuron connectivity in the brain. **(a)** Using the in-degree vs. out-degree scatterplot, we can divide the intrinsic rich club neurons into three distinct categories: broadcasters, integrators, and large balanced neurons. Comparing the prevalence of **(b)** neurotransmitters and **(c)** intrinsic superclasses (optic lobe intrinsic, visual projection, visual centrifugal, and central brain intrinsic) of all intrinsic neurons, rich club neurons, integrators, and broadcasters. **(d)** Examples of rich club neurons in these three categories. **(e)** Applying the information flow model from Schlegel et al. 2021 (28, 48), we determined the percentile rank distributions of rich club, integrator, and broadcaster neuron populations from all inputs to the brain (above), as well as to specific modalities (**Figure S4d**). **(f)** Average percentile rank of rich club, integrator, and broadcaster neurons for different modalities. Across all modalities, rich club neurons are closer than average to sensory inputs.

**Figure 5.**
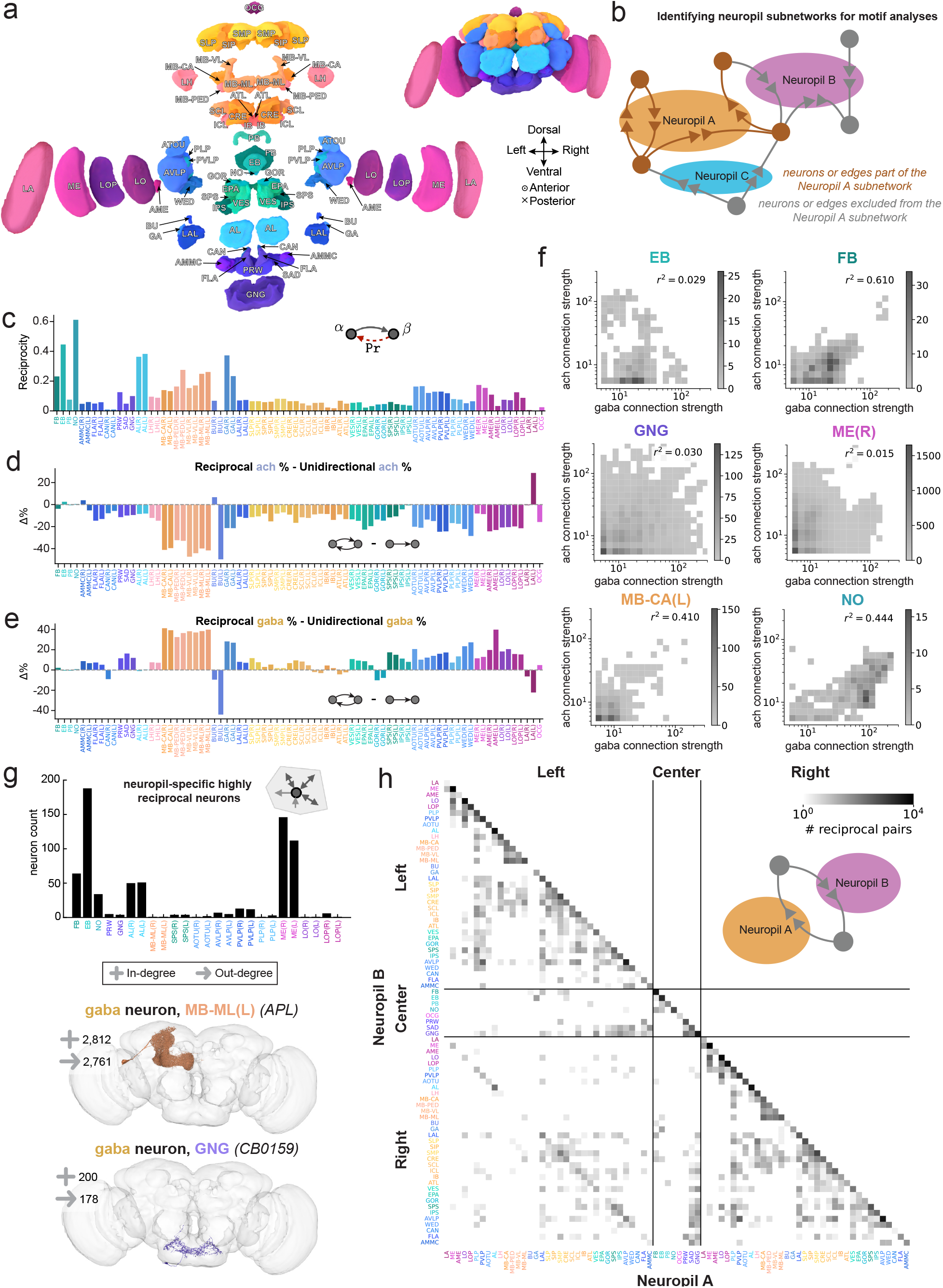
Neuropil-specific differences in connectivity. **(a)** An exploded view of the brain showing the brain regions, or neuropils, that the FlyWire dataset is divided into. Each synapse is assigned to a neuropil based on synapse location. **(b)** A schematic showing how neuropil subnetworks are identified for motif analyses. With the standard threshold of 5 synapses per edge applied, all connections composed of synapses within the neuropil of interest (Neuropil A) are treated as edges of the Neuropil A subnetwork. All neurons reached by this set of edges are included in the subnetwork. However connections composed of synapses outside of Neuropil A are not included, even if those connections involve neurons included in the subnetwork. **(c)** The reciprocity within each neuropil subnetwork. Differences in the percentage of **(d)** cholinergic and **(e)** GABAergic edges between reciprocal and unidirectional connections, across different neuropils. Refer to **Figure S6** for the absolute percentages. **(f)** Heatmaps showing the relationship between excitatory and inhibitory connection strengths in reciprocal connections in different brain regions. **(g)** Assessing the number of large (rich club), highly reciprocal neurons which span specific neuropils: making most of their incoming and outgoing connections within a single neuropil and also having a high reciprocal degree. Examples of neurons which meet these criteria are shown. **(h)** Map of the total number of reciprocal pairs between different neuropils. Examples of such pairs are shown in **Figure S7e**.

When compared to the population of all neurons, rich club neurons are less likely to be cholinergic and more likely to be GABAergic (**Figures 4b, S4a**). Integrator neurons are even less likely to be cholinergic (49%), and include a large fraction of dopaminergic neurons, suggesting that these neurons may be engaged during learning. In contrast, broadcaster neurons are predominantly cholinergic (75%). Central brain neurons are dramatically over-represented in the rich club, while optic lobe intrinsic neurons are under-represented (**Figures 4c, S4b**). Many integrators are either central brain intrinsic neurons or visual projection neurons. In contrast, few broadcasters are intrinsic to the central brain—many are visual centrifugal neurons or optic lobe intrinsic neurons. These include a large number of Mi1 and Tm3 neurons, excitatory cells in the medullae (ME) known to play key roles in the motion detection circuit (41, 49, 50). Most neurons are restricted to a single hemisphere—just 11% of neurons have inputs in both hemispheres and 11% have outputs in both hemispheres (**Figure S4c**)(28). In comparison, rich club neurons are more likely to have inputs or outputs spanning both hemispheres: 18% and 17%, respectively. This is more common for integrator neurons (23%) than it is for broadcaster neurons (16%).

### Rich club neurons are closer on average to sensory inputs

To assess the distance of the rich club neurons from sensory inputs, we employed a probabilistic information flow model to determine the relative distance of each neuron (in hops) from a set of seed neurons (**Methods**) (28, 48). The model was run with different sets of seed neurons, each corresponding to a specific set of sensory neurons (olfactory, gustatory, etc.), as well as on the complete set of all sensory inputs, giving us the distance from each neuron in the dataset to each sensory modal-ity. We excluded the visual photoreceptors from this analysis **(Methods)**. Ranking these distances and normalizing returned the percentile rank of each neuron with respect to each modality. Neurons with percentile rank less than 50% are closer than average to the given sensory input, while neurons with percentile rank greater than 50% are farther.

The rich club neurons have a mean percentile rank of 44% relative to the set of all sensory inputs (**Figure 4e**). Integrators have a mean percentile rank of 43%, while broadcasters have a mean percentile rank of 53%. Integrator neurons are closest, with many having a percentile rank of less than 10%. The distribution of broadcasters is bifurcated, with one peak closer to inputs and another peak far from inputs. Examining the ranks with respect to individual sensory modalities, we find that rich club neurons are again closer than average to each modality (**Figures 4f, S4d**). Broadcasters tend to be closer to single sensory inputs than they are to the set of all inputs. This is likely because ranking from a seed population of all inputs will rank integrators before many broadcasters. In contrast, when looking at a single modality, neurons which are predominantly connected to a different modality will be farther than average.

We examine the distance of neurons to multiple sensory modalities by plotting the percentile rank of neurons with respect to one modality against the percentile rank of neurons of another modality (**Figure S4e**). Broadcaster and integrator neurons are scattered throughout these distributions, but tend to be closer than average to multiple sensory inputs. These rich club neurons may be a fruitful starting point when searching for neurons to characterize experimentally. In particular, integrator and broadcaster neurons which are low in rank relative to multiple sensory modalities may be good candidate sites of multi-sensory integration and information propagation.

### Differences in connectivity across brain regions

The fly brain consists of a large number of distinct anatomical brain regions, or neuropils (51). The FlyWire connectome has been segmented into 78 neuropils (**Figure 5a**), each with different average connection strengths (28). To understand information flow between neuropils, we employed a fractional weighting method accounting for each neuron’s projections to and from every neuropil (**Methods**)(28). From these, we computed for each neuropil the relative fraction of internal, external incoming, and external outgoing connection weights (**Figure S5a-b**). These fractions reflect, respectively, the net number of connections within, being received, and being sent from each neuropil.

We find significant differences in these fractions across brain regions: the ellipsoid body (EB) and fan-shaped body (FB) of the central complex have the highest fraction of internal connections, while in other regions, such as the compartments of the mushroom body (MB), the majority of connections are external (**Figure S5b**). Some regions such as the lateral horn (LH) send more external connections than they receive, while others such as the lobula plate (LOP) receive more external connections than they send. The fraction of internal connection weights is not correlated with neuropil size: while large neuropils such as the anterior and posterior ventrolateral protocerebra (AVLP and PVLP) have significant fractions of internal weights, they do not rank the highest. We note that under this classification, internal weights include any neurons with endings outside the brain, such as sensory, ascending, and descending neurons. This likely accounts for the high fraction of internal weights in regions such as the medullae (ME), which receive inputs from R7 and R8 photoreceptors, and the gnathal ganglia (GNG), which connects with large numbers of both ascending and descending neurons. Across the brain, 52% of all connection weights can be classified as internal. Comparing the putative neurotransmitters of the neurons contributing connection weights, we see that internal connections are more likely than external ones to be inhibitory (GABAergic or glutamatergic) (**Figure S5c**). We also see differences in neurotransmitter composition across brain regions (**Figure S5d**).

### Prevalence and neurotransmitter composition of reciprocal connections differ across neuropils

To perform motif analyses within each neuropil, we first identified a subnetwork for each neuropil which treats all connections made within that neuropil as edges and includes all neurons connected to these edges (**Figures 5b, S6a**). Different neuropil subnetworks differ notably in both connection strength and density (**Figure S6b**). We computed the reciprocity in each neuropil subnetwork (**Figures 5c, S6c**). Neuropils with particularly high reciprocity probabilities include those in the central complex (FB, EB, and NO) and the two antennal lobes (AL). The relative number of reciprocal connections (reciprocity normalized by neuropil connection density) is high in the mushroom bodies (MB) and medullae (ME) (**Figure S6b**). Note that for these motif analyses, the results for small neuropils such as the cantles (CAN), bulbs (BU), galls (GA), accessory medullae (AME), and ocellar ganglion (OCG) are less interpretable due to the small number of samples.

In most neuropils, as in the whole brain, reciprocal connections are stronger than unidirectional connections, though the ratio of average strengths varies across neuropils (**Figure S6d**). Exceptions include the protocerebral bridge (PB), mushroom body calyces (MB-CA), and bulbs (BU), which have stronger unidirectional connections than reciprocal connections. Comparing the relative prevalence of each neurotransmitter in reciprocal and unidirectional connections, we again see differences between neuropils (**Figures 5d-e, S6d-h**). While reciprocal connections in most neuropils contain fewer cholinergic edges and more GABAergic edges than unidirectional connections, there are notable exceptions, such as in the neuropils of the central complex (FB, EB, PB, and NO). In the compartments of the mushroom body (MB) we find especially large differences in neurotransmitter composition between unidirectional and reciprocal connections. Comparing the strengths of the edges of reciprocal excitatory-inhibitory (acetylcholine-GABA) connections within neuropil subnetworks, we observe that E-I connection strengths are more strongly correlated in some neuropils (such as the FB and NO) than in others (**Figures 5f, S7a-b**). These correlations do not appear to be dependent on neuropil size (**Figure S7c**).

### Identifying neuropil-specific reciprocal neurons

We performed a comprehensive search for intrinsic highly reciprocal rich club neurons that make the majority of their connections within a single neuropil, and found 1,863 neurons that meet these criteria (**Figure 5g**). These *neuropil-specific highly reciprocal neurons* (NSRNs) are predominantly inhibitory: 54% are GABAergic and another 10% are glutamatergic (**Figure S7d**). In some neuropils, such as the antennal lobes (AL), medullae (MB), and ellipsoid body (EB), there are many NSRNs, while in other neuropils, such as the superior posterior slopes (SPS) and posteriorlateral protocerebra (PLP), there exist only a handful of such neurons.

Some NSRNs, like the APL neurons in the MB (52, 53), CT1 neurons in the LO (41, 54, 55), or antennal lobe local neurons (ALLNs) (56, 57), have been previously characterized as providing global feedback inhibition in different regions. These neurons tend to be highly branched, with individual processes making reciprocal connections with different feedforward neurons. Some have been shown to have compartmentalized activity, raising the possibility of local computation within these neurons (58–60). Many of the NSRNs identified here have yet to be characterized. They may play similar roles in other circuits— for instance, it is likely that some of the NSRNs found in the AVLP provide feedback to the auditory circuits which span this brain region (61).

### Identifying inter-neuropil reciprocal connections

While many reciprocal connections occur within single neuropils, 12.1% of all reciprocal pairs are formed by connections made by synapses in two neuropils (**Methods**). We mapped the reciprocal connections that exist between the 78 neuropils (**Figure 5h**). The diagonal terms consist of the intra-neuropil reciprocal connections described above (**Figure 5b-c**), while the off-diagonal terms reflect the number of reciprocal pairs which connect across neuropils. Examples of such neuron pairs are shown in **Figure S7e**.

From the map, we see that reciprocal connections exist between many neuropil pairs. The compartments of the mushroom body (MB) are linked by many reciprocal connections, while the neuropils of the SEZ, including the GNG, SAD, and PRW, form a connected block. Strong reciprocal connectivity also occurs across the midline. For instance, there is strong reciprocal connectivity between the two antenna lobes (AL(L) and AL(R)). Neuropils close to the midline, such as SMP, SPS, and IPS, tend to have many cross-hemispheric reciprocal connections. There also exist reciprocal connections which span from one edge of the central brain to the other, such as those between AOTU(L) and AOTU(R) and between LAL(L) and LAL(R). The prevalence of such inter-neuropil reciprocal connections demonstrates that the recurrent motifs we observe in the brain are not limited to local connections—they can also exist at large spatial scales.

Additional insight can be gleaned by comparing the map of reciprocal connections to the projectome matrix of all neurons in the brain (Dorkenwald et al., Figure 4 (28)). Comparing the two maps, we can identify regions which are connected by many neurons, but have disproportionately few reciprocal connections. For instance, neuropils SLP and SIP are connected to the FB in the projectome, but share no reciprocal connections. Similarly, the LA boasts many neurons but very few reciprocal connections.

Examining ach-gaba reciprocal connections, we can identify deviations from symmetry that represent a net imbalance of excitatory-inhibitory reciprocal connections (**Figure S7f**). For example, between the LO and PVLP, all ach-gaba reciprocal connections share the same directionality: the ach connections are in the LO and the gaba connections are in the PVLP.

### Three-node motifs differ across neuropils in their prevalence and strength

We computed the prevalence of three-node motifs in each neuropil subnetwork, and compared the motif frequencies to ER and CFG random null models constructed for each sub-network (**Figures 6a, S8a**). Across most neuropils, we observed the same trend as we do across the entire brain: an under-representation of feedforward motifs (#1-3) and an overrepresentation of complex motifs (**Figure 6b**). However, there are notable differences between neuropils. In the cantles (CAN), epaulettes (EPA), and gorgets (GOR), for example, the frequency of 3-node motifs was closer to that expected in a CFG null model, while in other neuropils like the ellipsoid body (EB), complex motifs are highly over-represented (**Figure 6b**).

**Figure 6.**
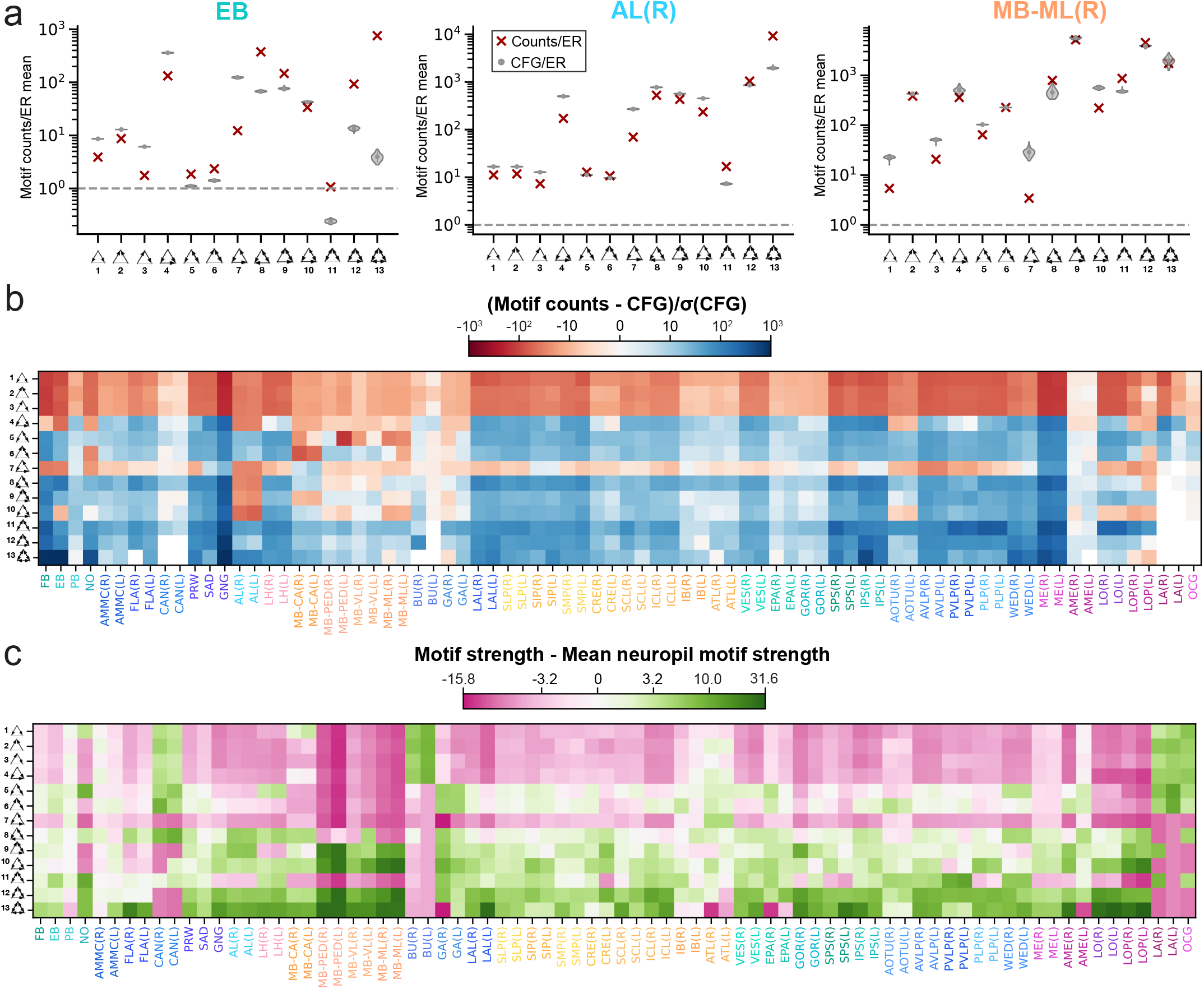
Differences in three-node motifs across neuropils. **(a)** Three-node motif distributions for three example neuropils: the EB, AL(R), and MB-ML(R). The frequency of each motif relative to that in an ER null model is plotted to the right, together with the average motif frequencies of 100 CFG models (gray violin plots). Further examples of other neuropils available in **Figure S8a. (b)** Motif frequencies for the 3-node motifs across all 78 neuropil subnetworks, normalized by their respective CFG null models. **(c)** Average strengths of edges participating in 3-node motifs in the different neuropil subnetworks relative to the average 3-node motif strength in each subnetwork. Refer to **Figure S8b** for average strengths relative to average neuropil subnetwork edge strength.

Feedforward loops (motif #4) are over-represented in most neuropils, excepting in the fan-shaped body (FB), ellipsoid body (EB), noduli (NO), and mushroom body compartments (MB). This suggests a relative under-representation of both feedforward excitatory and feedforward inhibitory circuits in these brain regions. 3-unicycles (motif #7), an indirect feed-back inhibition circuit, are over-represented across the whole brain (**Figure 3c**) but are under-represented in most neuropils. The notable exceptions, the medullae (ME) and gnathal ganglia (GNG), are very large neuropils and have many sensory inputs. The over-representation of 3-unicycles in the ME implies the existence of localized cyclic structures within the early visual circuitry. Interestingly, this motif is also over-represented in the zebrafish oculomotor circuit (7). Motifs #7-10 are underrepresented in the antennal lobes (AL), perhaps a result of the small number of unidirectional edges in these regions. The most highly connected motifs (#12-13) are particularly over-represented in the ellipsoid body (EB) and fan-shaped body (FB), consistent with their high reciprocity.

In most neuropils, we find that edges participating in under-represented motifs are also weaker on average than edges participating in over-represented motifs (**Figure 6c**). We also observe that in most neuropil subnetworks, edges participating in 3-node motifs are stronger than the average subnetwork edge (**Figure S8b**). This is broadly consistent with the whole-brain 3-node motif strength results. A notable exception is in the laminae (LA), where feedforward connections are strong despite being under-represented.

## Discussion

Here, we have provided a broad overview of the network properties of the *Drosophila* brain, laying the groundwork for identifying neurons and circuit motifs of biological interest and for modeling of particular circuits. In addition to the topology of the neural network, we have taken advantage of spatial information (innervation in different neuropils), neuron class distinctions (sensory versus descending, for example), cell type labels, and neurotransmitter predictions to better contextualize and interpret the network features we uncovered. We compared the statistics of the fly connectome to other wiring diagrams, carried out a comprehensive brain-wide search for 2- and 3-node connectivity motifs, identified highly connected broadcaster and integrator neurons, and identified differences in connectivity in different brain regions. The complete FlyWire dataset is freely available online via Codex (Connectome Data Explorer: codex.flywire.ai), along with interactive lists of the neurons discussed in this work. These data will allow researchers to profile neurons by their connectivity features and identify key neurons within their circuits or brain regions of interest, a useful resource for hypothesis generation or model development. Experimentally examining highly connected neurons, such as the attractors, repellers, integrators, broadcasters, and NSRNs identified here, may also prove fruitful for linking circuit-level findings with broader activity patterns. Our results reveal that despite its sparsity, the neurons of the brain form a robust and highly interconnected network. This network is not predominantly feedforward, with over-represented reciprocal and recurrent motifs which can span multiple brain regions. Additionally, different brain regions in the fly differ in their network properties.

An understanding of how the whole-brain network shapes brain function is particularly important in light of recent experimental findings. A common approach in modern experimental neuroscience is to use anatomical wiring diagrams to generate circuit-level hypotheses, and to test these hypotheses by imaging and perturbing single cells or cell types. However, recent whole-brain imaging experiments, both in the fly (62–64) and in other species (65–72), have revealed brain-wide activity patterns related to both sensory processing (of individual modalities) and simple behaviors (like locomotion). To fully understand distributed computations and information flow in the brain, we must consider interactions not just at the scale of tens of neurons, but at the brain scale. Availability of network statistics at the scale of brain regions, coupled with the broad mesoscale connectivity between brain regions (28), will enable hypothesis generation at the whole-brain scale. Different neuropils serve different functions, and our work now highlights how these different functions are subserved by differences in connection strength, internal connectivity, motif frequency, and neurotransmitter composition. For example, the central complex (neuropils FB, EB, PB, and NO), which has persistent activity associated with an internal representation of heading (73– 78), contains some of the most reciprocal brain regions and has a large number of internal connections. Examination of other neuropil subnetworks may help us generate hypotheses regarding the function of less well-studied neuropils.

In this work, we comprehensively explored 2-node and 3-node motifs, and highlighted several large-scale connectivity patterns by exploring broadcaster (few-to-many), integrator (many-to-few), and highly reciprocal neurons. There remains, of course, a space of larger network motifs to explore. We have integrated the network motif search and visualization tool Vimo (79) into Codex, which allows users to query the FlyWire connectome for any network motif of interest.

### Limitations

The availability of neurotransmitter predictions greatly enhanced our ability to interpret the circuit motifs we found in the connectome. However, while these predictions are 94% accurate when compared to a set of ground truth neurons, there are cases where the predicted neurotransmitter does not align with the known transmitter. In this iteration of the dataset, we manually corrected the Kenyon cells to be cholinergic (**Methods**). There may exist other populations of neurons which are likewise systematically mis-identified, but which currently lack ground truth neurotransmitter information. When interpreting results on the network scale, we must keep this error rate in mind. Also, monoamines beyond dopamine, octopamine, and seratonin are not accounted for in these predictions. More details on the neurotransmitter predictions are discussed in Eckstein *et al*. (33). In this work, we assume that neurons in the fly obey Dale’s law—each releasing only one neurotransmitter. However, there are several known examples of co-transmission in *Drosophila* (80–83). How widespread neurotransmitter co-transmission is remains unclear.

It should also be noted that the synaptic connectome does not provide a complete picture of information flow in the brain. We currently do not have a complete map of gap junctions in the fly, and the extent to which extrasynaptic communication (via non-synaptic release of amines or neuropeptides) shapes neural activity in the *Drosophila* brain remains an open question (84– 86).

We also acknowledge that some of the statistics presented here, particularly those metrics dependent on network topology, such as neuron degree or reciprocity and motif frequencies, may be sensitive to our choice of synapse threshold. While connectomes in *C. elegans* have been proofread to the level of individual synapses (2, 19, 25, 44), it is not feasible to manually proofread every synapse in larger connectomics datasets (27, 28, 38). We must therefore rely on automated synapse detection algorithms with a non-negligible error rate (32). Not all synapses are successfully attached to neurons, and this completion rate varies across animals and brain regions (24, 28, 38). To avoid false positive connections, we applied a threshold on the number of synapses a connection between neurons must have. While some of these low synapse number connections may be spurious, it is also likely that a significant number of these weak connections are real and reliable across individuals, as has been found when comparing multiple individuals in *C. elegans* (25). In this work, we employed a consistent and conservative threshold of five synapses per connection between neurons, and demonstrated that our qualitative conclusions are not dependent on this threshold. We therefore analyzed a sparser network of high-confidence connections, containing 2.6 million connections instead of 14.7 million un-thresholded connections (**Table S2**). It is likely that the fly brain is even more strongly interconnected than the results here indicate.

Local circuit motifs are often inferred to be feedforward or feedback connections, with different theorized roles. While we are able to make such inferences on the population level, it can be difficult to place local circuits in the context of global directionality from sensory input to motor output. In shallow networks such as in *C. elegans*, the directionality of the wiring diagram from sensory input to motor output is clear. However, the larger the network becomes, the more difficult it becomes to establish directionality from sensory input to motor output. In this work, we employed an information flow method to rank the neurons by an effective difference from various sensory modalities (28). Ultimately, however, directionality of information flow in particular circuits, especially those in regions of the brain far from sensory inputs or motor outputs, must be determined through functional activity experiments and modeling.

### The rich club compensates for anatomical bottlenecks

The anatomy of the fly brain suggests several potential network bottlenecks: one between left hemisphere and right hemisphere and one between the central brain and optic lobes. Only 12% of neurons cross hemispheres and 6% of neurons cross between the central brain and optic lobes (28, 29). Despite these bottlenecks, the brain is robustly interconnected with short path lengths. The large rich club regime in the fly brain may explain these short path lengths. When compared to the average neuron in the brain, rich club neurons are more likely to contain synapses in both hemispheres, and are also more likely to connect the optic lobes to the central brain. The broad reach of these rich club neurons also keeps path lengths short across these bottlenecks. In mesoscale functional connectome work in the human brain, it has similarly been proposed that rich-club hubs act to keep path lengths short (87, 88). Future functional imaging experiments in the fly focusing on the population of rich club neurons may shed light on whether this this is the case at neuron-scale.

We may also expect the ascending and descending neurons which form a bottleneck between the brain and the ventral nerve cord (VNC) will also be part of a rich club of the central nervous system. Many ascending and descending neurons appear to have high degrees when examined either within the brain or within the VNC. While a wiring diagram of the VNC is now available (89), we await the completion of a complete CNS connectome to determine whether the ascending and descending neurons are members of the rich club.

### Comparing connectomes across animals

Comparing network properties across wiring diagrams from different species has the potential to uncover global properties of brain organization. We make several such comparisons in **Table 2**, and have commented on other comparisons throughout the text. The similarities in reciprocity and clustering coefficient across animals, which vary dramatically in both size and connection density, hint at the possibility that some features of circuit architecture may be broadly conserved across biological nervous systems. Comparisons of metrics which are dependent on network topology, however, such as neuron degree or reciprocity and motif frequencies, must be interpreted with care due to differences in proofreading and data resolution. While connectomes in *C. elegans* have been proofread to the level of individual synapses (2, 19, 25, 44), in larger connectomics datasets individual synapses are not proofread and instead a threshold on synapses per connection is applied to filter out spurious connections (24, 27, 28, 38). Threshold choice impacts topological metrics, which treat all edges as equivalent. Applying the same threshold across datasets does not resolve this conundrum, as a given number of synapses per connection may have different biological implications across species. It has also been observed, both in this work and in past studies, that different parts of the brain of the fly differ in their connectivity properties (38, 42). It is likely that the same is true in larger, more complex brains as well, meaning that statistics derived from partial wiring diagrams may not be representative.

It has been demonstrated in *C. elegans* that there is substantial variability in the connectomes of individuals of the same species (25). Comparisons between the FlyWire connectome and hemibrain wiring diagram have already revealed interesting similarities and differences between individual flies, as outlined in our companion paper (29), but more datasets will be needed before we fully understand the amount of variability between individuals in *Drosophila*. The same is expected to be true for zebrafish and mouse connectomes. More whole-brain connectomes are on the horizon, both in *Drosophila* and in other species (90). The network analysis of the fly brain presented here will be a valuable baseline for comparison, both to the connectomes of other *Drosophila* individuals and to the connectomes of other species. As the efficiency of electron microscopy and neural reconstruction continue to increase, it will become possible to better understand which features of these networks are common and which are species- or individual-specific. Such comparative connectomics studies within a single species may shed light on brain development, stereotypy, and learning, while future studies across multiple organisms may elucidate principles of brain evolution, organization, and computation.

## Methods

### Dataset

The FlyWire connectome is the reconstruction of a 7-day-old adult female *Drosophila melanogaster*, genotype [iso] w1118 x [iso] Canton-S G1 (30). The EM images were aligned and neurons were automatically reconstructed using deep learning and computer vision methods, then proofread by the community (27, 28). Neuron cell types and community labels were also attached to these data (29, 91). All analyses presented in this paper were performed on the *v*630 Snapshot of the Fly-Wire dataset. The *v*630 snapshot contains 127,978 neurons and 2,613,129 thresholded connections, the central brain of the fly was fully proofread, with the optic lobes *∼*80% complete. Most of the neurons missing from the *v*630 Snapshot were photoreceptors, and we do not expect that the addition of these neurons would significantly change our whole-brain network results. At time of publication, the most up-to-date version of the FlyWire dataset is the *v*783 Snapshot, containing 139,255 neurons, 2,701,601 thresholded connections, and completed optic lobes. Both data snapshots are available at Codex (Connectome Data Explorer): codex.flywire.ai.

### Synaptic connections and thresholding

Synapses were detected algorithmically (31, 32), with each synapse receiving a confidence score. We then removed synapses if (1) either the pre- or postsynaptic location of the synapse was not assigned to a segment, or (2) the synapse had a confidence score of less than 50. We then set a threshold of 5 synapses per connection between neurons for most of our analyses to reduce the impact of spurious connections. This threshold is also consistent across our companion papers on the FlyWire connectome (28, 29). We employed a threshold because manual proofreading of the FlyWire dataset did not extend to individual synapses (28). Thresholding connections by synapse number was previously implemented in the hemibrain connectome, with similar rationale (38). We acknowledge that this is a conservative threshold and is likely to result in an undercounting of true connections. We assessed key statistics as a function the threshold to ensure that our qualitative observations hold over a range of threshold choices (**Figure S1b-c**).

### Assignment of neurotransmitters to neurons

The neurotransmitter at each synapse was predicted directly from the EM images using a trained convolutional neural network with per-synapse accuracy of 87% (28, 33). The algorithm returns a 1 *×* 6 probability vector containing the odds that a given synapse is each of the six primary neurotransmitters in *Drosophila*: ach, gaba, glut, da, oct, or ser. We then averaged these probabilities across all of a neuron’s outgoing synapses, under the assumption that each neuron expresses a single outgoing neurotransmitter, to obtain a 1 *×* 6 probability vector representing the odds that a given neuron expresses a given neurotransmitter. We then assigned the highest-probability neurotransmitter as the putative neurotransmitter for that neuron. The per-neuron accuracy is 94%.

In cases where the highest probability is *p*_1_ *<* 0.2 and the difference between the top two probabilities *p*_1_ *− p*_2_ *<* 0.1, we classified the neuron as having an uncertain neurotransmitter. In the *∼*1600 Kenyon cells, where the neurotransmitter of a neuron is known to be acetylcholine but the algorithm often returned erroneous predictions, the neurotransmitter prediction associated with that neuron was overwritten by the known neurotransmitter.

### Cell classifications and labels

84% of neurons are *intrinsic* to the brain, meaning that their projections are fully contained in the brain volume (28). Central brain neurons are fully contained in the central brain, while optic lobe intrinsic neurons are fully contained in the optic lobes. Visual projection neurons have inputs in the optic lobes and outputs in the central brain. Visual centrifugal neurons have inputs in the central brain and outputs in the optic lobe. Sensory neurons are those which are entering the brain from the periphery, and are divided into classes by modality. Refer to our companion paper, Schlegel et al., for more details on classification criteria (29). We also employed annotation labels contributed by the FlyWire community (28).

### Definitions of in-degree, out-degree, total degree and reciprocal degree

For a given neuron *i*, the in-degree 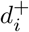 is the number of incoming synaptic partners the neuron has and the out-degree 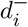 is the number of outgoing synaptic partners the neuron has. The total degree of a neuron *i* is the sum of in-degree and out-degree:

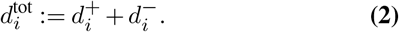

The reciprocal degree 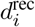 is the number of partners a given neuron form reciprocal connections with. Since each reciprocal connection consists of two edges, we can determine the fraction of reciprocal inputs and outputs as 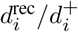 and 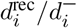, respectively (**Figure S3c**).

### Definitions of connection probability, reciprocity and clustering coefficient

Given the observed wiring diagram as a simple (no self-edges) directed graph *G*(*V, E*), the “connection probability” or “density” is the probability that, given an ordered pair of neurons *α* and *β*, a directed connection exists from one to the other:

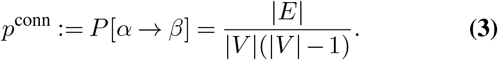

The reciprocity is the probability that, given a pair of neurons which are connected *α* to *β*, there exists a returning *β* to *α* connection:

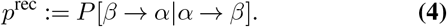

The (global) clustering coefficient is the probability that for three neurons *α, β* and *γ*, given that neurons *α* and *β* are connected and neurons *α* and *γ* are connected (regardless of directionality), neurons *β* and *γ* are connected:

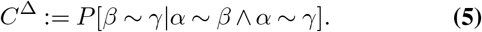

We computed these metrics both across the whole brain and within brain region (neuropil) subnetworks.

We also systematically quantified the occurrence of distinct directed 3-node motifs within the network, ensuring that duplicates are eliminated: any subgraph involving three unique nodes is counted only once in our analysis. To compute the expected prevalence of specific neurotransmitter motifs (**Figures 2c, 3d-e**) we multiplied the relevant neurotransmitter probabilities for the motif of interest, under the assumption the neurons connect independent of neurotransmitter. We then compared this expectation to the true frequency of motifs with these neurotransmitter combinations.

### ER and CFG null models

We probed different statistics of the wiring diagram *G*(*V, E*) by comparing them with the statistics of various null models. The simplest null model we employed was a directed version of the Erdő s–Rényi model (ER) *G*(*V, p*), where all edges are drawn independently at random, and the connection probability *p* is set such that the expected number of edges in the ER model equals that observed in the wiring diagram (47). For any nodes *i, j ∈ V*, the connection probability is constant:

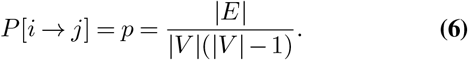

Since reciprocal edges in the wiring diagram are over-represented when compared to a standard Erdő s-Rényi (ER) model, we adopted a generalized Erdős–Rényi model (gER), which preserves the expected number of reciprocal edges. The gER model *G*(*V, p*^uni^, *p*^bi^) has two parameters, uni-directional connection probability *p*^uni^ and bi-directional connection probability *p*^bi^, both of which are set to match the wiring diagram. To do this, we defined the sets of unidirectional and bidirectional edges as:

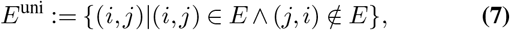

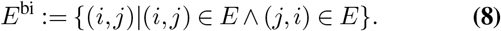

For any nodes *i* and *j*:

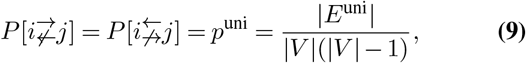

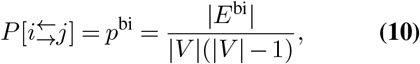

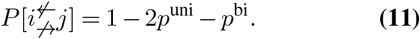

All edges between unordered node pairs were drawn independently and at random.

In line with previous work (7, 63), we also employed a directed configuration model (CFG), (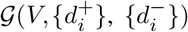), which preserves degree sequences during random rewiring. We sampled 1,000 random graphs uniformly from a configuration space of graphs with the same degree sequences as the observed graph by applying the switch-and-hold algorithm (92), where we randomly select two edges in each iteration and swap their target endpoints under the condition that doing so does not introduce self-loops or multiple edges (switch), or else keep them unchanged (hold).

### Computing pairwise distances between neuronal arbors

To determine the connection probability distribution as a function of distance between neurons, we first had to distill the available spatial information into a handful of points. This was the only practical way to enable distance comparisons between all neurons—a total of 14 billion pairs.

For each neuron, we defined two coordinates based on the location of their incoming and outgoing synapses. We computed the average 3D position of all of the neuron’s incoming synapses to approximate the position of the neuron’s dendritic arbor, and did the same to approximate the position of the neuron’s axonal arbor. We then computed for all neuron pairs the pairwise distances between the axonal arbor of neuron *A* and the dendritic arbor of neuron *B*. Binning by distance and comparing the number of true connections to the number of neuron pairs allowed us to compute connection probability as a function of space (**Figure S1d**).

### Spatial null model

Informed by the distribution of connection probability as a function of distance, we constructed a spatial null model with two zones of probability—a “close” zone (0 to 50 microns) where connections are possible with a relatively high probability (*p*_*close*_ = 0.00418) and a “distant” zone (more than 50 microns) where connections occur with lower probability (*p*_*distant*_ = 0.00418) (**Figure S1e**). The probabilities in these two zones were derived from the real network.

### Spectral analysis

Given a strongly connected graph *G*(*V, E*) and its 0-1 adja-cency matrix 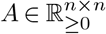, where *A*_*ij*_ indicates the existence of a connection from neuron *j* to neuron *i*, one can construct an irreducible Markov chain on the strongly connected graph with a transition matrix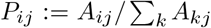 giving the transition probability from *j* to *i*. The Perron-Frobenius theorem guarantees that *P* has a unique positive right eigenvector *π* with eigenvalue 1, and therefore that *π* is the stationary distribution of the Markov chain. We constructed such a transition matrix for the connectome and determined the eigenvector *π*.

We also defined a “reverse” Markov chain with a transition matrix 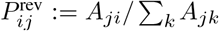giving the transition probability from *j* to *i* also has a unique positive right eigenvector *π* ^rev^ with eigenvalue 1. **Figures** S1f and S1g show the stationary distribution of forward and reversed Markov chains, respectively.

The normalized symmetric Laplacian of the Markov chain *P* is

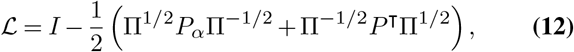

where Π:= `Diag`(*π*) and *I* is the identity matrix. Similarly, we defined *L*^rev^ for the reverse Markov chain. The eigen-spectra of *L*^rev^ and *L*^rev^ are shown in **Figures S1f** and **S1g**, respectively. The gaps between eigenvalues indicate the conductance properties of the graph.

### Finding rich club neurons

We employed the standard rich club formulation to quantify the rich club effect (11). The rich club coefficient Φ(*k*) at a given degree value (*k*), with all nodes with degree *< k* pruned, is the number of existing connections in the surviving subnetwork divided by the total possible connections in the surviving subnet-work:

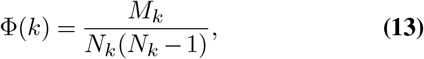

where *N*_*k*_ neurons in the network with degree *≥ k* and *M*_*k*_ is the number of connections between such neurons.

To control for the fact that high-degree nodes have a higher probability of connecting to each other by chance, we normalized the rich club coefficient to the average rich club value of 100 samples from a CFG null model **(Figure 1h)**:

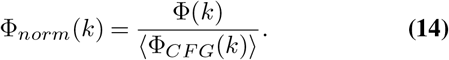

The standard method of determining the rich club threshold is to look for values of *k* for which Φ_*norm*_(*k*) *>* 1 + *nσ*, where *σ* is the standard deviation of Φ_*CF G*_(*k*) and *n* is chosen arbitrarily (18). However, since the standard deviation from our samples is extremely small near the bump in relative rich club coefficient, we chose instead to define the onset threshold of the rich club as Φ_*norm*_(*k*) *>* 1.01 (1% denser than the CFG random networks). We computed the rich club coefficient in three different ways, by sweeping by total degree (Figure 1h), in-degree, and out-degree (Figure S2c), progressively moving from small to large values. As we observed, when the total degrees of the remaining nodes surpass 37, the network becomes denser compared to randomized networks. Once the minimal total degree reaches 93, the network becomes as sparse as the randomized counterpart. Therefore, we classified neurons with total degrees above 37 as “rich club” neurons because they exhibit denser interconnections when considered as a subnetwork. In terms of in-degree, the range for denser-than-random connectivity is between 10 and 54. Considering out-degree alone did not reveal any specific onset or offset threshold for rich club behavior, as the subnetwork always remains sparser than random.

### Definitions of broadcaster neurons, integrator neurons, and neuropil-specific recurrent neurons

To identify broadcaster neurons, we filtered the intrinsic rich club neurons (*d*^tot^ *>* 37) for those which had an out-degree was at least 5 times higher than their in-degree:

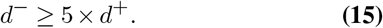

Similarly, we identified integrator neurons by filtering the intrinsic rich club neurons for those which had an in-degree was at least 5 times higher than their out-degree:

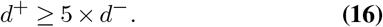

Rich club neurons which did not fall into either category were defined as “large balanced” neurons. This analysis was limited to intrinsic neurons–those which have all of their inputs and outputs within the brain–to avoid spurious identification of afferent or efferent neurons as broadcasters or integrators.

When identifying large recurrent neuropil-specific neurons (**Figure 5g**) we applied the following criteria. First, the neurons were intrinsic and met the rich club criteria. Second, at least 50% of the neuron’s incoming connections were contained within the subnetwork of a single neuropil. Third, at least 50% of the neuron’s outgoing connections were contained within the same neuropil.

### Neuron ranking

We employed a probabilistic connectome flow model previously published in Schlegel et al. 2021 to determine the ranking of neurons relative to various sensory neuron populations (28, 48). This method ignores the sign of connections. Starting from a set of user-defined seed neurons, the model traverses the wiring diagram probabilistically: in each iteration the chance that a neuron is added to the traversed set increases linearly with the fractions of synapses it is receiving from neurons already in the traversed set. When this likelihood reaches 30%, the neuron is guaranteed to be added to the traversed set. The process is then repeated until the entire network graph has been traversed. The iteration in which a neuron was added corresponds to the distance in hops it was from the seed neurons. For each set of seed neurons, the model was run 10,000 times. The distance used to determine the rank of any given neuron was the average iteration in which it was added to the traversed set.

We ran this model using the following subsets of sensory neurons as seeds: olfactory receptor neurons, gustatory receptor neurons, mechanosensory Johnston’s Organ neurons, head and neck bristle mechanosensory neurons, thermosensory neurons, hygrosensory neurons, visual projection neurons, visual photoreceptors, ocellar photoreceptors and ascending neurons. We also ran the model using the set of all of the input neurons as seed neurons. All neurons in the brain were then ranked by their traversal distance from each set of starting neurons, and this ranking was normalized to return a percentile rank.

### Determining information flow between neuropils

To determine the contributions a single neuron makes to information flow between neuropils, we first applied two simplifying assumptions: (1) that information flow through the neuron can be approximated by the fraction of synapses in a given region and (2) that inputs and outputs can be treated independently. Employing these two assumptions we constructed a matrix representing the projections of a single neuron between neuropils. The fractional inputs of a given neuron are a 1 *× N* vector containing the fraction of incoming synapses the neuron has in each of the *N* neuropils, and the fractional outputs are a similar vector containing the fraction of outgoing synapses in each of the *N* neuropils. We multiplied these vectors against each other to generate the *N × N* matrix of the neuron’s fractional weights, with a total weight of one. Summing these matrices across all neurons produced a matrix of neuropil-to-neuropil connectivity, or projectome (see Figure 4 of Dorkenwald et al., 2023) (28).

From the neuropil-to-neuropil connectivity matrix we determined the total weight of internal connections—those within a given neuropil—by identifying the neurons which contribute to the diagonal of the matrix. We likewise determined the weight external connections—either incoming to the neuropil or outgoing from the neuropil—by looking at the off-diagonals. These data were used to construct the analyses in **Figure S6a-c**.

### Identifying neuropil subnetworks

Most of the neurons in the *Drosophila* brain have soma at the surface of the brain. Therefore, they cannot be associated to neuropils (brain regions) based on their soma locations. Synapses, however, can be associated with neuropils. To perform motif analyses at the level of individual neuropils, we identified neuropil subnetworks based on the the connections made by the synapses contained within each neuropil volume. All connections within the neuropil of interest are taken as edges of this subnetwork, and all neurons connected to these edges are included (**Figure 5b**). The number of neurons associated with each neuropil subnetwork is plotted in **Figure S6d**. Note that if two neurons both in a given neuropil subnetwork share a connection which occurs in a different neuropil, that connection is not included as an edge in the given subnetwork.

### Identifying inter-neuropil reciprocal pairs

We constructed a map of reciprocal connections between neuropils in the form of a triangular matrix with the neuropils as axes. For clarity, here we will refer to a unidirectional connection as an edge. A reciprocal connection contains two opposing edges. While some edges are composed of synapses in multiple neuropils, the majority of edges are composed of synapses in a single neuropil after thresholding. We therefore applied a winner-take-all approach to assigning edges to neuropils.

Given two recipocally connected neurons X and Y, let us call the edge from X to Y Edge 1, and the edge from Y to X Edge 2. If the synapses that form Edge 1 are in Neuropil A, and the synapses that form Edge 2 are in Neuropil B, then we assign this reciprocal pair to the Neuropil A to Neuropil B square of the matrix. This was done for all reciprocal pairs, with each reciprocal pair is counted as 1 in the matrix. Note that this means that a given neuron can be represented multiple times if it has multiple reciprocal partners.

### Data availability

The FlyWire data is available online via Codex (Connectome Data Explorer): codex.flywire.ai. Neuron annotations, neurotransmitter information, and compact data downloads are available via Codex, along with neuron lists generated in this work, including neurons participating in 2-node and selected 3-node motifs, rich club neurons, broadcaster and integrator neurons, and neuropil-specific reciprocal neurons.

### Software availability

The analyses presented in this paper were performed in Python with the numpy and graph-tool (93) packages, and in MATLAB (standard toolboxes). Software written for this publication is available at Github (github.com/murthylab/flywire-network-analysis). Some 3D renders were generated in Cinema4D.

### AUTHOR CONTRIBUTIONS

AL, RY, and SD analyzed the data. SD, PS, and AM curated the data and made it available for download. AM developed software and data analysis tools. AM and ARS built the Codex online platform. SCY, CEM, MC, KE, and PS trained and managed Flywire proofreaders. ASB, NE, GSXEJ and JF provided neurotransmitter information. CEM and ARS provided Flywire community support and training. AL, RY, SD, and ARS generated figure panels. AL, RY, and MM wrote the manuscript with feedback from the other authors. MM supervised the project.

## ACKNOWLEDGEMENTS

We thank Sebastian Seung, Davi Bock and the members of the Murthy, Seung, Jefferis, and Clandinin labs for their advice on the project and comments on the manuscript. AL was supported by the NSF through the Center for the Physics of Biological Function (PHY-1734030). GSXEJ was supported by Wellcome Trust Collaborative Awards 203261/Z/16/Z and 220343/Z/20/Z, the Neuronex2 award (MRC_MC_EX_MR/ T046279/1), and the MRC (MC-U105188491). MM was supported by NIH BRAIN Initiative grants RF1 MH117815, RF1 MH129268 and U24 NS126935. We also acknowledge support from the Princeton Neuroscience Institute and assistance from Google.

## Supplement Supplemental figures

**Figure S1.**
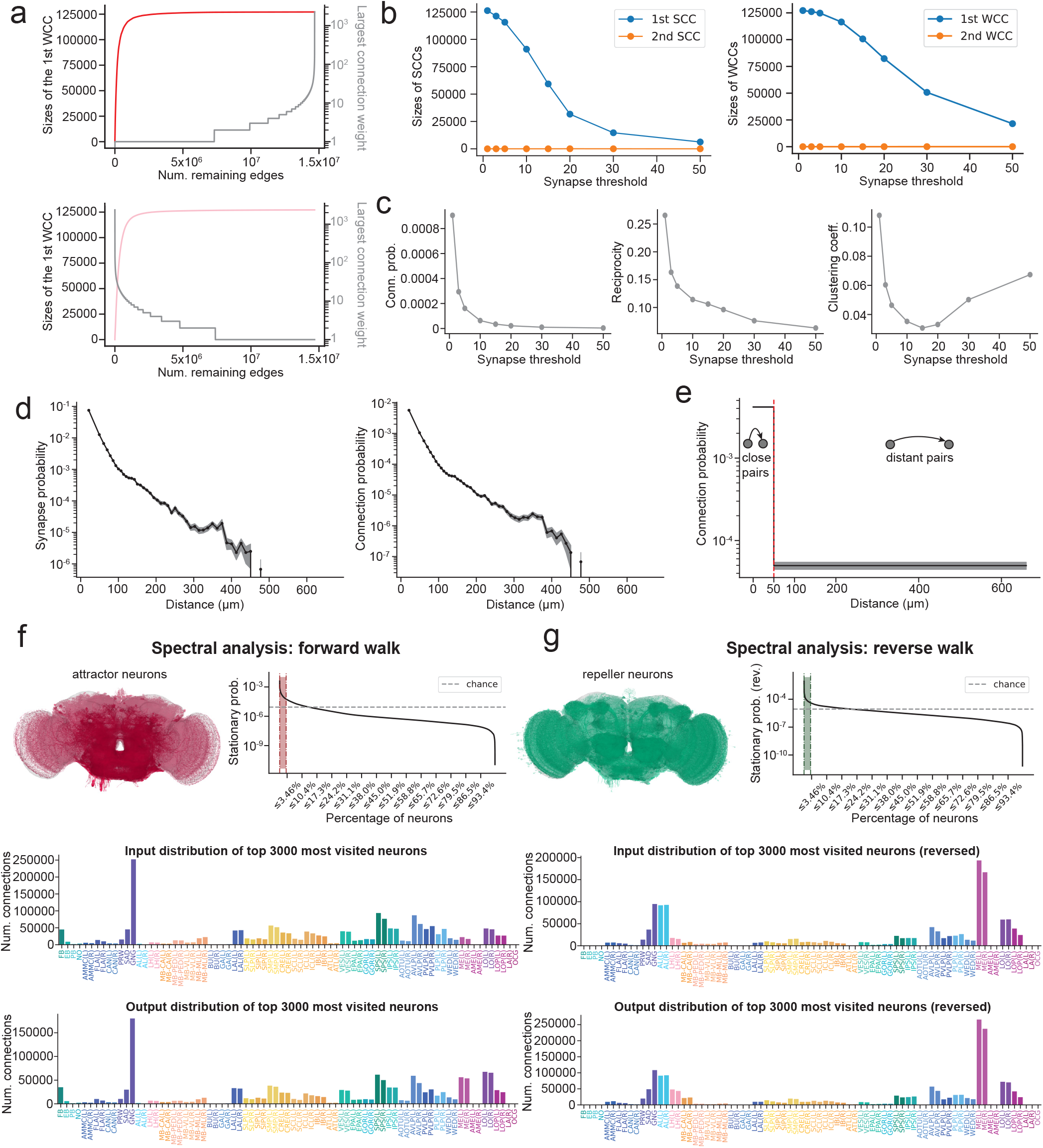
Supplement for Figure 1. The effects of edge percolation on the size of the largest WCC when **(a)** large connections are removed first and when **(b)** small connections are removed first. **(c)** The sizes of the first two SCCs as a function of the synapse threshold. **(d)** Synapse probability (left) and connection probability (right) as a function of the average distance between neuronal arbors. Plots are of a drawn from a subsample of 700 million pairs (5% of the total 14 billion pairs). **(e)** The probability of random connection of the two-zone spatial null model, with one close regime with high connection probability and a distant regime with low connection probability. Spectral analysis of the whole-brain network with **(f)** forward and **(g)** reverse walks. In each case, the stationary probability distributions are shown, as well as the distribution of neuropils in which the inputs and outputs of the top 3000 most visited neurons are located. Renders of the top 3% attractor (red) and repeller (green) neurons are also shown. The top 0.3% are rendered in darker colors.

**Figure S2.**
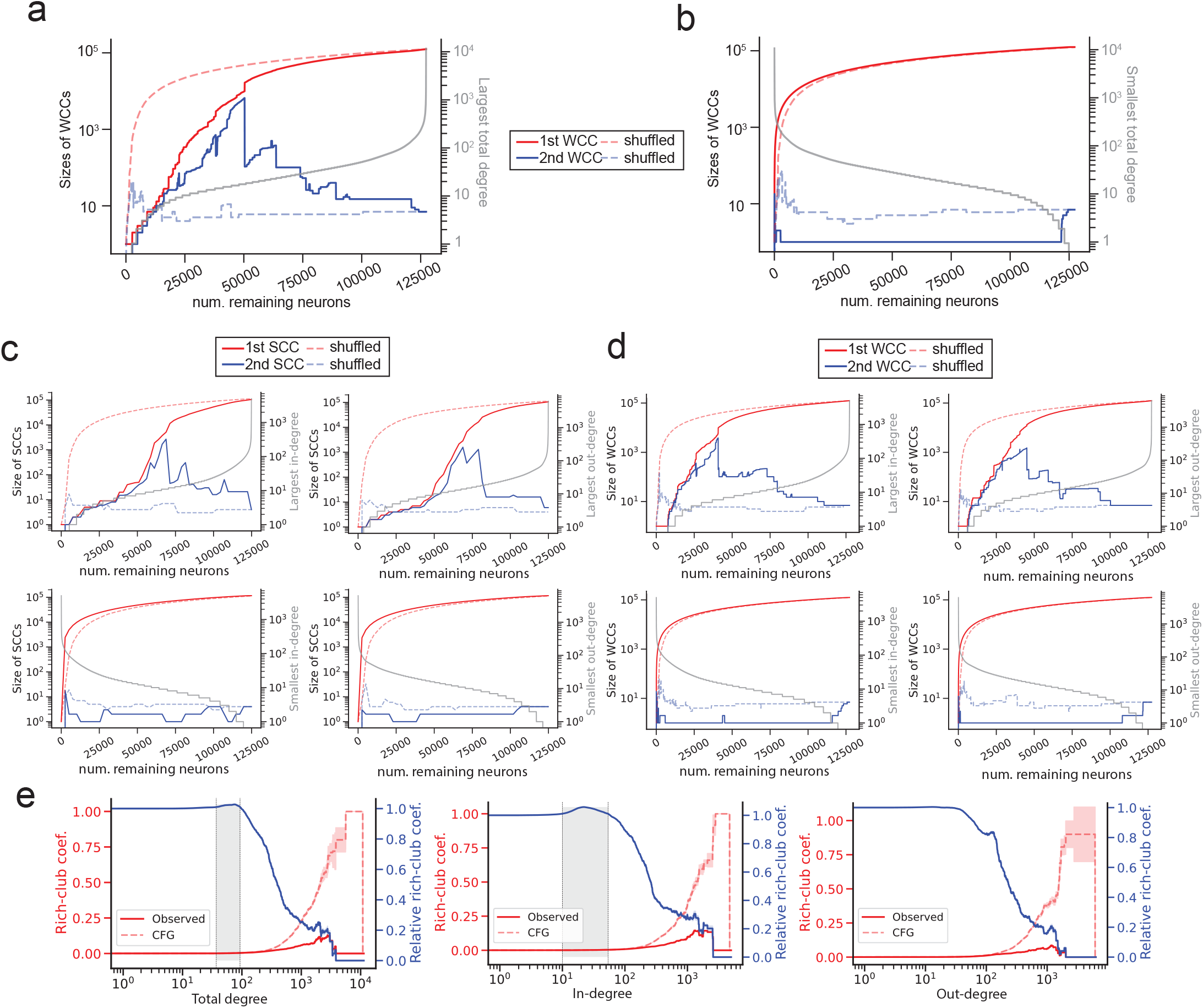
Additional supplement for Figure 1. **(a)** The sizes of the first two weakly connected components (WCCs) as nodes are removed by total degree (1 neuron per step). Removal of neurons starting with those with largest degree results in the brain splitting into two WCCs when neurons of approximately degree 50 start to be removed, a deviation from when neurons are removed in a random order (dotted lines). The largest surviving total degree as a function of the number of remaining nodes is plotted in gray. **(b)** Removal of neurons starting with those with smallest degree results in a single giant WCC until all neurons are removed. The smallest surviving total degree as a function of the number of remaining nodes is plotted in gray. **(c)** The sizes of the first two strongly connected components (SCCs) as nodes are removed by in-degree or out-degree (2500 neurons per step). Removal of neurons starting with those with largest in-degree (top left) or largest out-degree (top right) result in the brain splitting into two SCCs when neurons of approximately degree 50 start to be removed, a deviation from when neurons are removed in a random order (dotted lines). Removal of neurons starting with those with smallest in-degree (bottom left) or smallest out-degree (bottom right) results in a single giant SCC until all neurons are removed. **(d)** The sizes of the first two weakly connected components (WCCs) as nodes are removed by in-degree or out-degree (1 neuron per step). Removal of neurons starting with those with largest in-degree (top left) or largest out-degree (top right) result in the brain similarly splitting into two WCCs when neurons of approximately degree 50 start to be removed, a deviation from when neurons are removed in a random order (dotted lines). Removal of neurons starting with those with smallest in-degree (bottom left) or smallest out-degree (bottom right) results in a single giant WCC until all neurons are removed. **(e)** The rich club coefficient (red) as a function of total degree (left), in-degree (middle), and out-degree (right), compared to the predicted rich-club coefficient of a CFG null model (dotted red). The relative rich club coefficient is plotted in blue

**Table S1.**
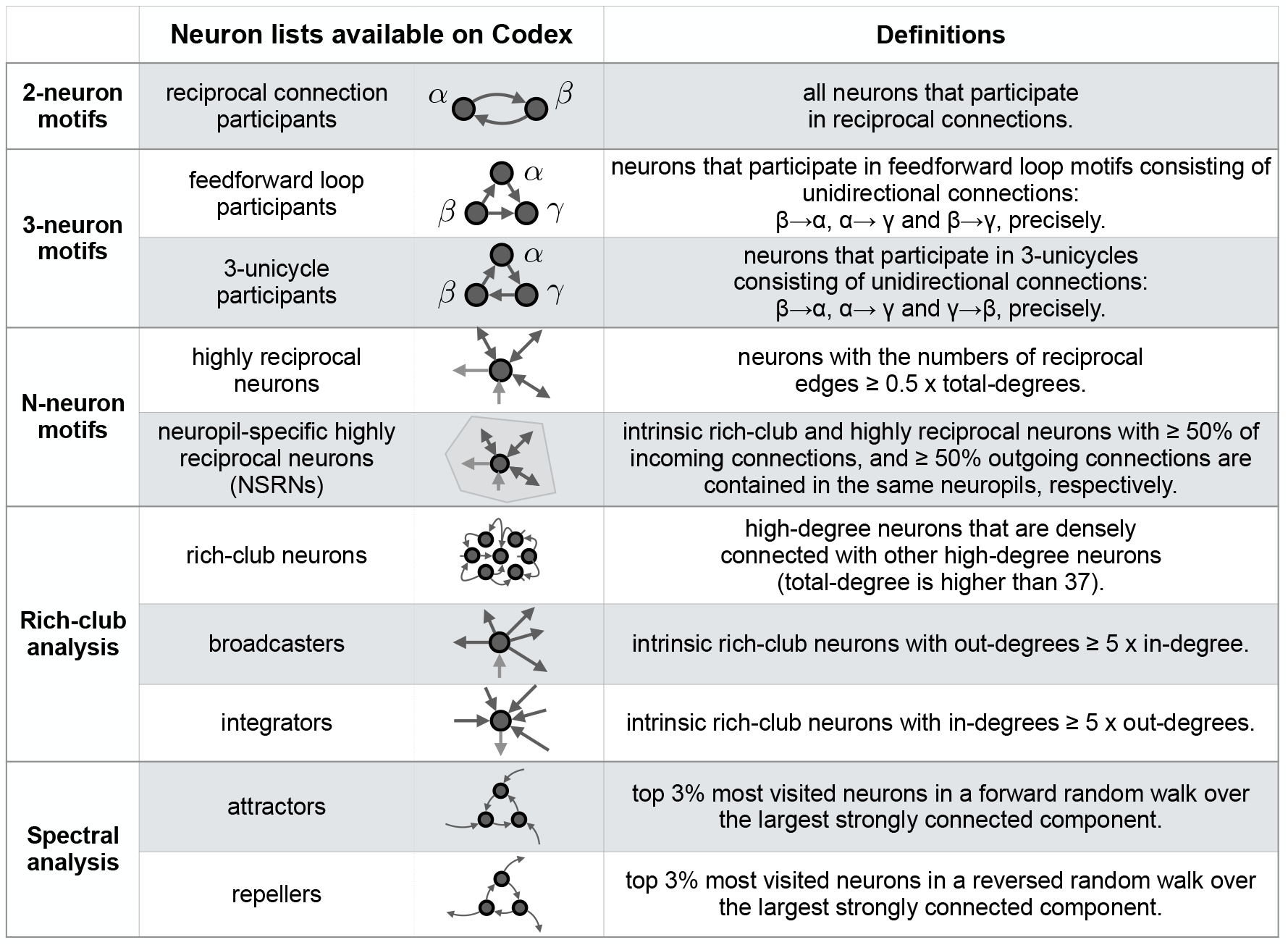
Supplement for Table 1. Definitions for all neuron populations identified in this paper.

|  | Neuron lists available on Codex |  | Definitions |
| --- | --- | --- | --- |
| **2-neuron motifs** | reciprocal connection participants |  | all neurons that participate in reciprocal connections. |
| **3-neuron motifs** | feedforward loop participants | | neurons that participate in feedforward loop motifs consisting of unidirectional connections: $\beta \rightarrow \alpha$, $\alpha \rightarrow \gamma$ and $\beta \rightarrow \gamma$, precisely. |
| | 3-unicycle participants | | neurons that participate in 3-unicycles consisting of unidirectional connections: $\beta \rightarrow \alpha$, $\alpha \rightarrow \gamma$ and $\gamma \rightarrow \beta$, precisely. |
| **N-neuron motifs** | highly reciprocal neurons | | neurons with the numbers of reciprocal edges $\geq 0.5 \times$ total-degrees. |
| | neuropil-specific highly reciprocal neurons (NSRNs) | | intrinsic rich-club and highly reciprocal neurons with $\geq 50\%$ of incoming connections, and $\geq 50\%$ outgoing connections are contained in the same neuropils, respectively. |
| **Rich-club analysis** | rich-club neurons |  | high-degree neurons that are densely connected with other high-degree neurons (total-degree is higher than 37). |
| | broadcasters | | intrinsic rich-club neurons with out-degrees $\geq 5 \times$ in-degree. |
| | integrators | | intrinsic rich-club neurons with in-degrees $\geq 5 \times$ out-degrees. |
| **Spectral analysis** | attractors |  | top 3% most visited neurons in a forward random walk over the largest strongly connected component. |
|  | repellers |  | top 3% most visited neurons in a reversed random walk over the largest strongly connected component. |

**Table S2.**
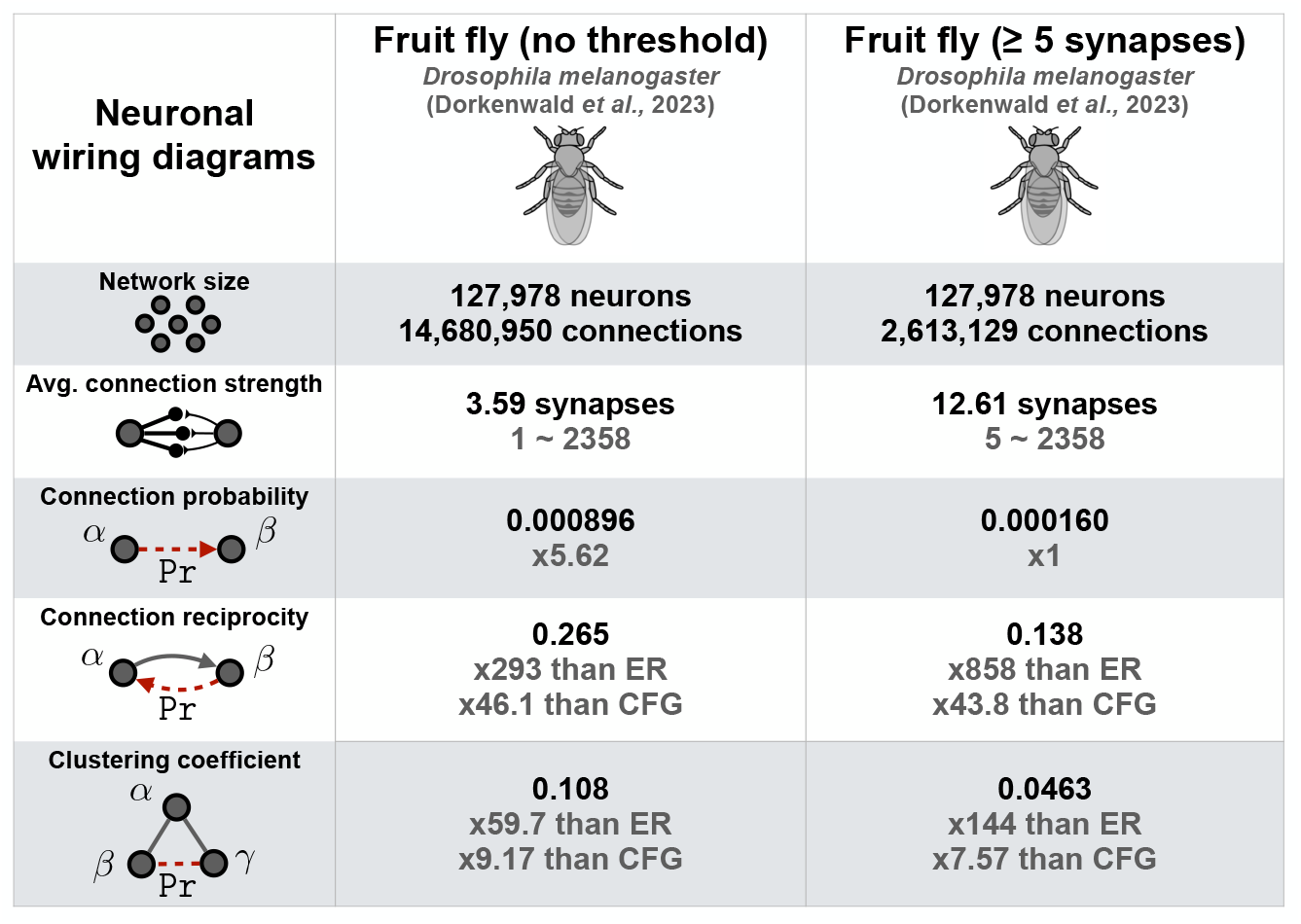
Supplement for Table 2. Network statistics of the fly connectome with no threshold on the number of synapses per connection (left) and a threshold of 5 synapses per connection.

| Neuronal wiring diagrams | Fruit fly (no threshold)<br>*Drosophila melanogaster*<br>(Dorkenwald et al., 2023) | Fruit fly ($\geq 5$ synapses)<br>*Drosophila melanogaster*<br>(Dorkenwald et al., 2023) |
| --- | --- | --- |
| Network size | **127,978 neurons**<br>**14,680,950 connections** | **127,978 neurons**<br>**2,613,129 connections** |
| Avg. connection strength | **3.59 synapses**<br>1 ~ 2358 | **12.61 synapses**<br>5 ~ 2358 |
| Connection probability | **0.000896**<br>x5.62 | **0.000160**<br>x1 |
| Connection reciprocity | **0.265**<br>x293 than ER<br>x46.1 than CFG | **0.138**<br>x858 than ER<br>x43.8 than CFG |
| Clustering coefficient | **0.108**<br>x59.7 than ER<br>x9.17 than CFG | **0.0463**<br>x144 than ER<br>x7.57 than CFG |

**Figure S3.**
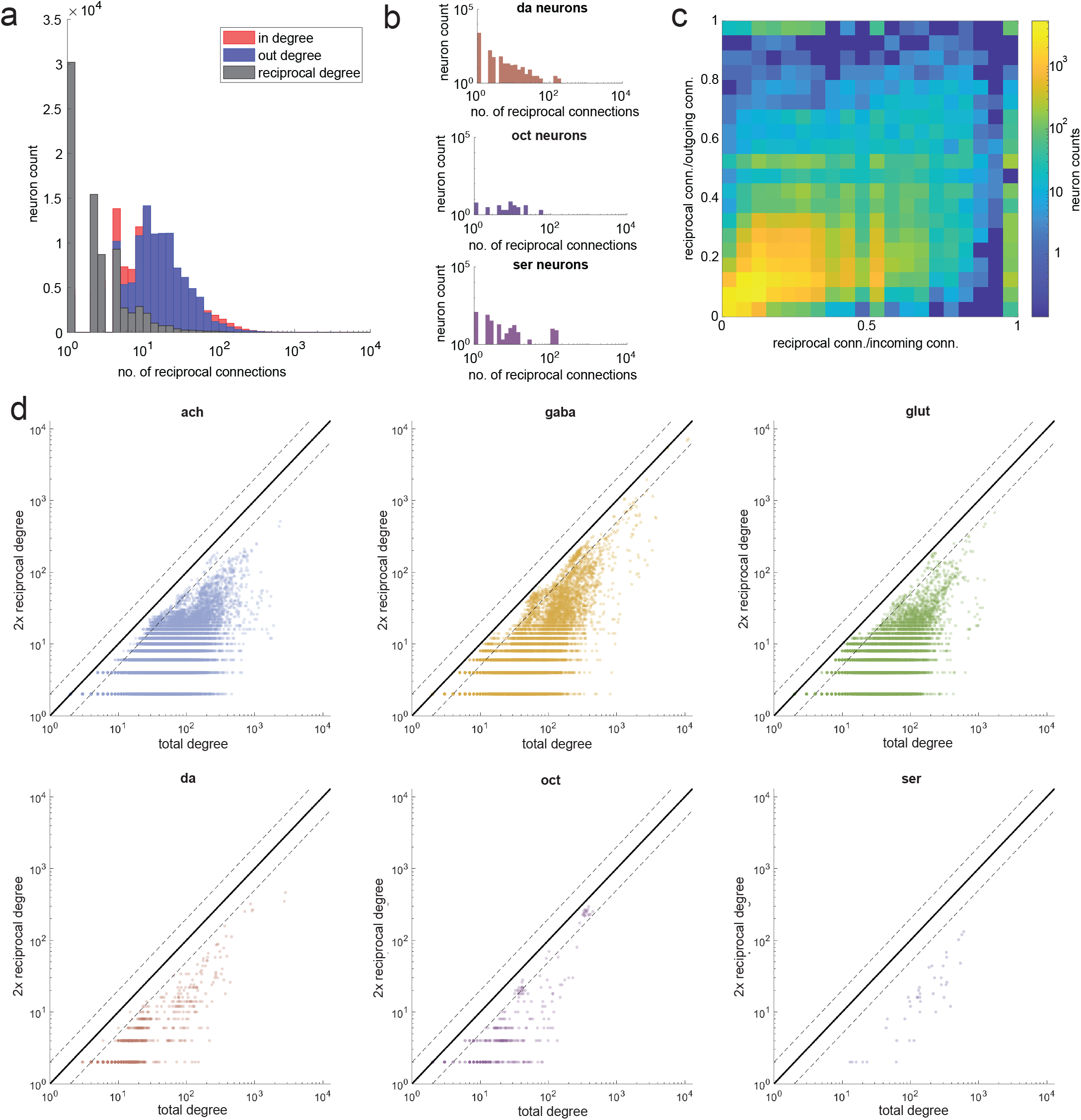
Supplement for Figure 2. **(a)** Distribution of reciprocal degree (gray) alongside distributions of in-degree (red) and out-degree (blue). **(b)** Distributions of reciprocal degree for glut, da, oct, and ser neurons. **(c)** Heatmap showing the fraction of reciprocal incoming connections versus the fraction of reciprocal outgoing connections. Dotted lines indicate a factor of 2 around the *x* = *y* line.

**Figure S4.**
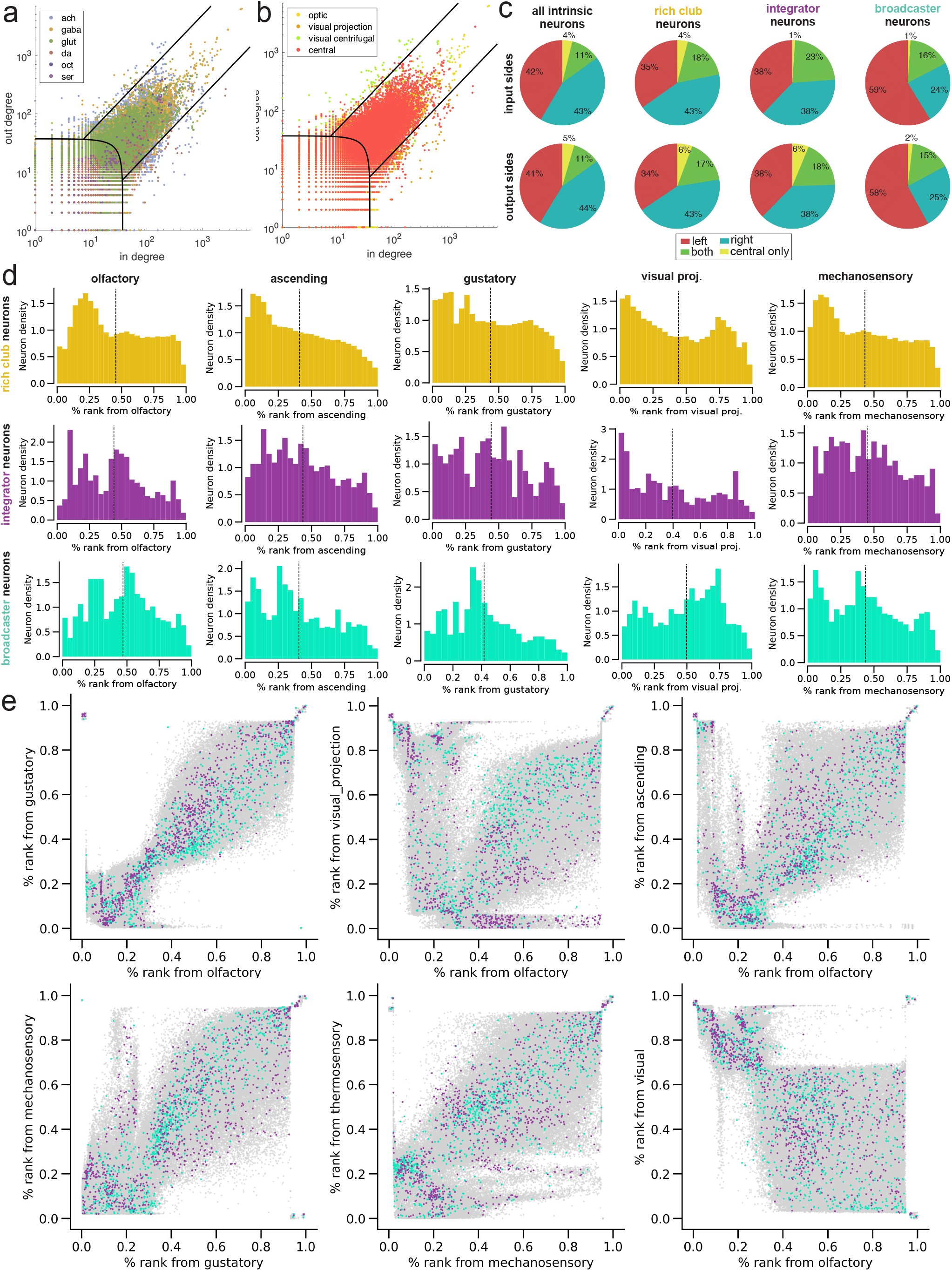
Supplement for Figure 4. In-degree vs. out-degree scatterplots showing broadcaster, rich balanced, and integrator regimes, with neurons plotted by **(a)** the putative neurotransmitter of each neuron and **(b)** the superclass of each neuron. **(c)** Comparing the input and output sides of all intrinsic neurons, rich club neurons, integrators, and broadcasters. The asymmetry in L/R percentages for broadcaster neurons is due to the large number of medulla-intrinsic broadcasters which connect with photoreceptors (Proofreading of photoreceptors was incomplete in Snapshot *v*630). **(d)** Percentile rank distributions of rich club, integrator, and broadcaster neuron populations from various input modalities. **(e)** Scatterplots of percentile rank from one sensory modality on each axis. Broadcaster neurons are highlighted in teal and integrator neurons are highlighted in purple.

**Figure S5.**
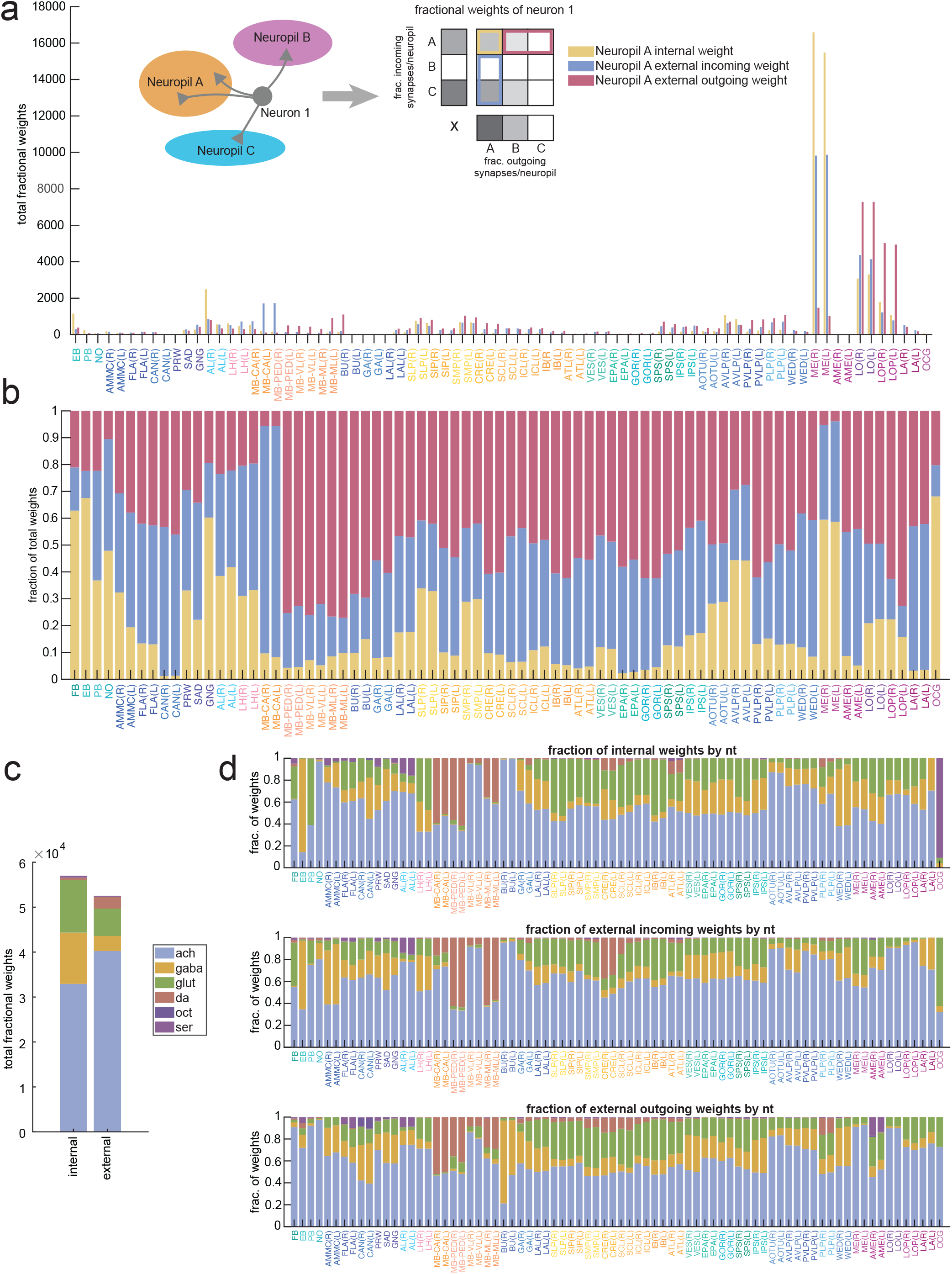
Internal and external connections across neuropils. **(a)** The number and **(b)** relative fraction of neuron weights in each neuropil making connections internal to that neuropil, external incoming connections, and external outgoing connections. Each neuron contributes a total weight of 1, computed based on the fraction of incoming and outgoing synapses the neuron has in each neuropil. **(c)** Comparing the neurotransmitter composition of all internal and all external neuron weights across the whole brain and **(d)** by neuropil.

**Figure S6.**
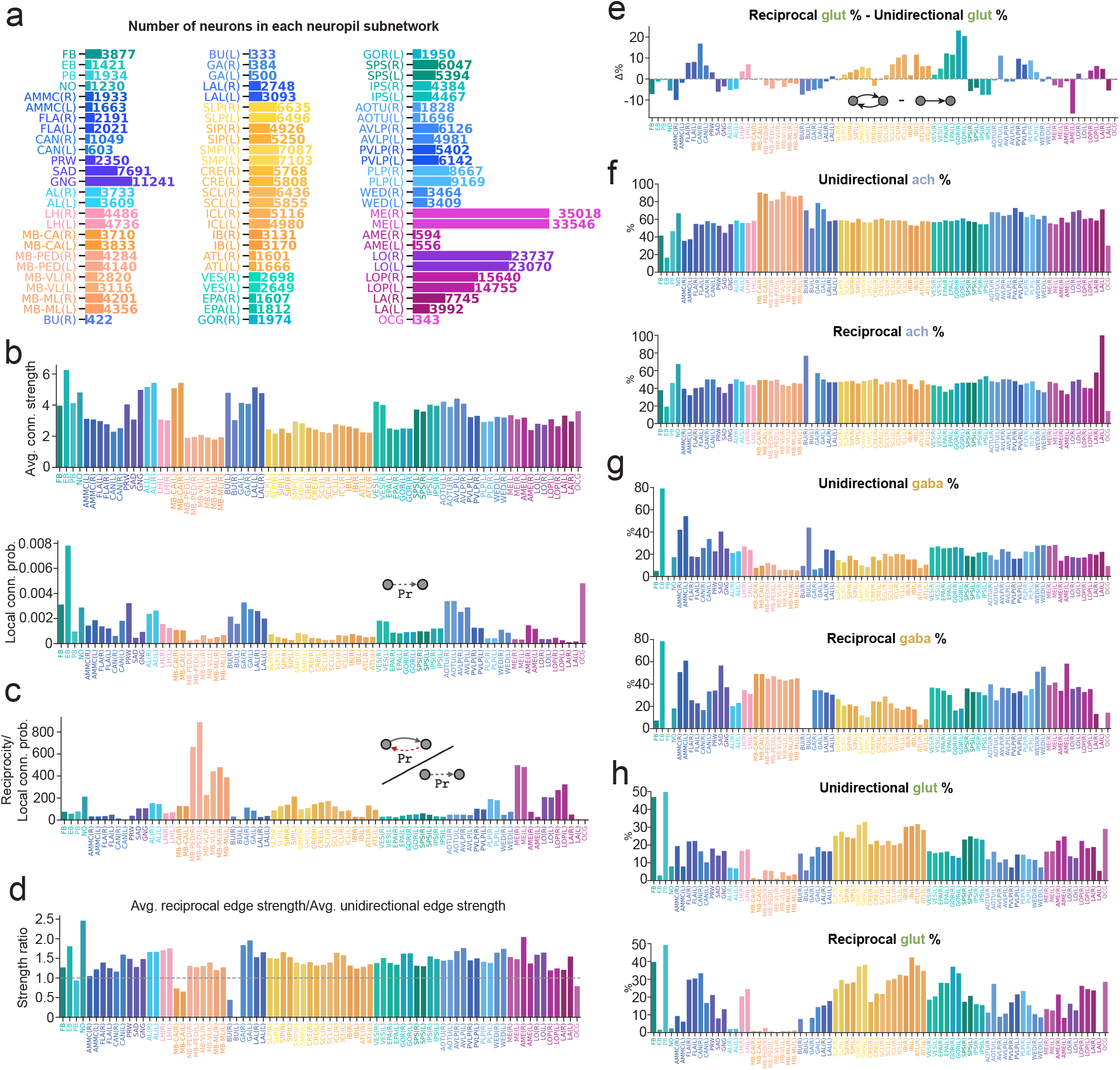
Supplement for Figure 5. **(a)** The number of neurons included in each neuropil subnetwork. **(b)** The average connection strength (no synapse threshold applied) of connections made in each neuropil (above), and the connection probability of each neuropil (below). **(c)** Reciprocity normalized by connection density for all 78 neuropils. **(d)** Average reciprocal connection strength normalized by average unidirectional connection strength in all neuropils. **(e)** The relative fraction of glutamatergic neurons participating in reciprocal and unidirectional connections. Absolute percentages of **(f)** acetylcholine, **(g)** GABA, and **(h)** glutamate occurrence in unidirectional and reciprocal connections within each neuropil subnetwork.

**Figure S7.**
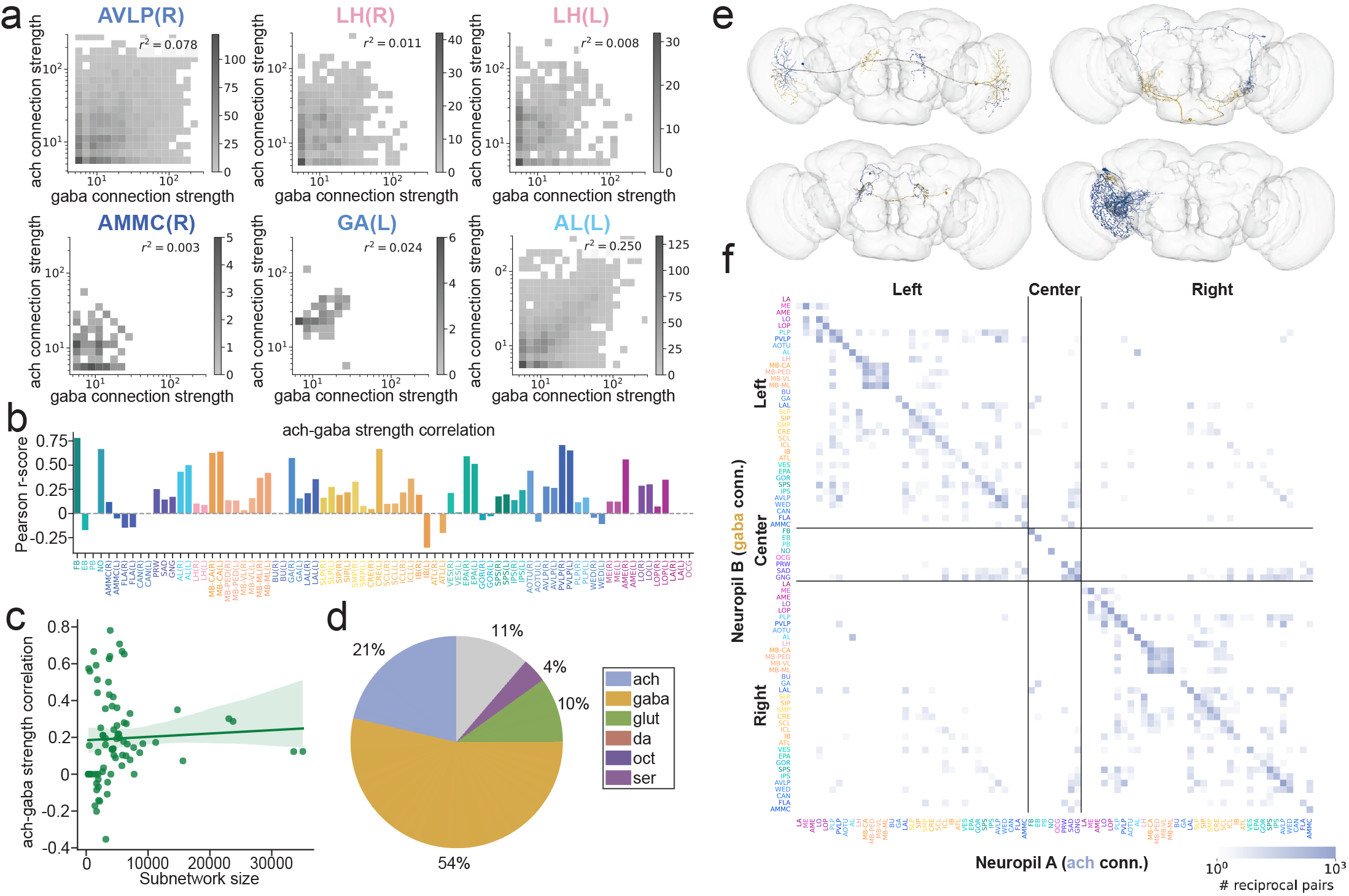
Additional supplement for Figure 5. **(a)** Heatmaps showing the relationship between excitatory (ach) and inhibitory (GABA) connection strengths in reciprocal connections in different brain regions. **(b)** Ach-gaba reciprocal connection strength correlations (Pearson r-score) for all neuropils. **(c)** These correlations do not appear to be correlated with neuropil subnetwork size. **(d)** The neurotransmitter composition of the population of neuropil-specific highly reciprocal neurons (NSRNs). **(e)** Examples of inter-neuropil reciprocal neuron pairs, one neuron in blue and one neuron in gold. **(f)** Map of the total number of ach-gaba reciprocal pairs between different neuropils.

**Figure S8.**
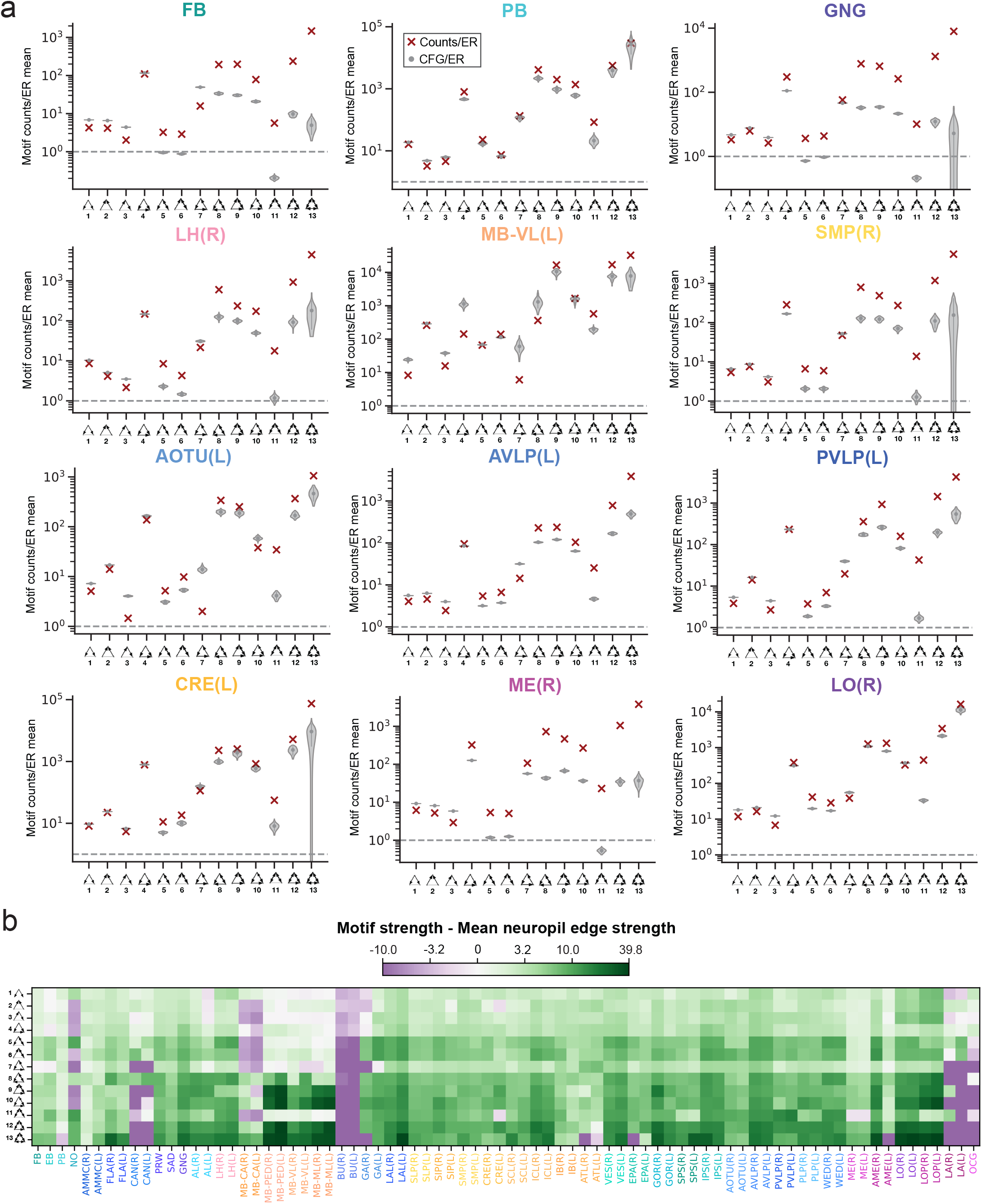
Supplement for Figure 6. **(a)** Three-node motif distributions for additional neuropils. The frequency of each motif relative to that in an ER null model is plotted to the right, together with the average motif frequencies of 100 CFG models (gray violin plots). **(b)** Average strengths of edges participating in 3-node motifs in the different neuropil subnetworks relative to the average edge strength in each subnetwork.

